# Interdependent androgen and glucocorticoid receptor signalling shapes prostate epithelial homeostasis

**DOI:** 10.64898/2026.07.09.737442

**Authors:** Qingshuang Cai, Tanisha Sooben, Anna-Isavella Rerra, Léa Bouhelier, Félicie Cottard, Daniela Rovito, Karim Essabri, Tao Ye, Emmanouela Epeslidou, Jia Wen Lim, Graciela Ruiz, Joe Rizk, Sirine Souali-Crespo, Rajesh Sahu, Emilia Calvano, Alicia Boule, Nandita Bodra, Hugo Rolando Vaca, Gilles Travé, Mariel Donzeau, Elodie Monsellier, Stefan Prekovic, Daniel Metzger, Isabelle M.L. Billas, Delphine Duteil

## Abstract

Androgen signalling is essential for prostate secretory functions and epithelial cell maintenance, yet how this signal is translated into a transcriptional output remains poorly understood. Androgens functions are primarily mediated by the androgen receptor (AR), which binds hormone response elements similar to those of other steroid receptors, including the glucocorticoid receptor (GR). Here we show that AR and GR are co-expressed in the prostate epithelium, and co-located within nuclear foci. Moreover, GR promotes the formation and dynamics of AR nuclear condensates, and heterodimerizes with AR in a ligand-binding domain-dependent manner. In addition, subtle variations within the hormone response element sequence direct the co-recruitment of AR and GR into activator or repressor complexes to activate or repress gene expression. Finally, prostate-specific deletion of GR in mice disrupts AR nuclear distribution, impairs AR-dependent gene networks involved in epithelial maintenance, and promotes the expression of genes involved in metabolism and cell cycle, resulting in altered tissue homeostasis. Thus, these findings identify GR as an integral component of the AR transcriptional machinery in epithelial cells of the healthy prostate, and reveal how shared cis-regulatory elements are interpreted to generate distinct transcriptional outcomes.

## Introduction

Androgens and glucocorticoids are steroid hormones that exert highly pleiotropic effects on mammalian physiology. Androgens primarily control development of sexual characteristics, behaviour, muscle mass and strength, as well as cell proliferation, whilst glucocorticoids are key regulators of immune response, circadian rhythm and energy metabolism. Increasing evidence indicates that androgen and glucocorticoid signalling pathways are highly interconnected. Early studies emphasized an antagonistic relationship, whereby androgens downregulate GR expression in various cell types^1–3^, and glucocorticoids attenuate androgen signaling^4^. Whether this crosstalk is limited to such mutual antagonism, or also extends to cooperative modes of regulation, has remained largely unexplored. The effects of androgens and glucocorticoids are mediated by the androgen receptor [AR (NR3C4)] and the glucocorticoid receptor [GR (NR3C1)], two ligand-dependent transcription factors, members of the nuclear receptor (NR) superfamily^5–10^. Both AR and GR share a modular domain architecture, comprising a large N-terminal intrinsically disordered region (NTD), a highly conserved DNA-binding domain (DBD), a flexible hinge region and a C-terminal ligand-binding domain (LBD)^11^. Phylogenetic studies have shown that AR and GR, together with the mineralocorticoid receptor [MR (NR3C2)] and the progesterone receptor [PR (NR3C3)], constitute the oxosteroid nuclear receptor subfamily^12^. These receptors are activated by natural 3-ketosteroid ligands and bind DNA predominantly as homodimers to a nearly identical response element composed of two 5’-RGAACA-3’ palindromic half-sites separated arranged as an inverted repeat separated by three nucleotides (IR3)^13–17^, hereafter referred to collectively as hormone response elements (HREs). In the classical model, ligand binding drives receptor nuclear translocation and engagement of these elements, historically designated androgen response elements (AREs) or glucocorticoid response elements (GREs), to activate target gene expression. In addition to activating transcription, both receptors can also mediate gene repression. GR represses transcription either indirectly by tethering to other transcription factors, such as AP-1 and NF-κB, to antagonize their activity (’transrepression’)^18^, or directly through negative response elements (nGREs)^19^. AR can likewise repress gene expression through poorly characterized transrepression mechanisms^20^, or *via* direct DNA binding in a context-dependent manner^21,22^.

Genome-wide studies in prostate cancer cell lines have shown that the AR and GR cistromes overlap by 25-45% depending on the cell type, and that approximately half of genes co-bound by both receptors are regulated by either one^1,23^. These findings indicate that AR and GR converge on common regulatory regions. However, whether such convergence reflects functional cooperation between the receptors, rather than redundant or competitive binding to similar DNA sequences, remains unresolved. A DNA-mediated AR-GR heterodimer has been proposed on the basis of a limited number of *in vitro* studies^24,25^, but whether such complexes form in cells and what functions they serve remain unknown. Moreover, current knowledge emerges almost exclusively from transformed, androgen-driven cancer cells, leaving the existence and physiological relevance of AR-GR cooperation in healthy tissues largely unexplored.

Like other transcription factors, AR and GR modulate gene expression in collaboration with co-regulatory proteins, whose recruitment is remodelled in a ligand-dependent manner^26–28^. Moreover, the intrinsically disordered NTDs of both receptors allow liquid-liquid phase separation (LLPS) and the formation of biomolecular condensates, which have been proposed to contribute to enhancer activity and to shape 3D genome architecture via the formation of super-enhancers^29–35^. Thus, the homotypic condensation of AR and GR has emerged as an important determinant of their transcriptional activity^29,31–33^. Nevertheless, whether the two receptors can co-assemble into shared condensates and how such higher-order organization influences transcriptional output remain unknown.

The prostate epithelium offers a unique physiological to address these questions, as it is one of the few tissues where AR and GR are co-expressed^36–38^. While AR has long been recognized as the master regulator of prostate epithelial identity^36,37^, the functional role of GR in the healthy prostate has received less attention^38^. Here, by integrating genome-wide analyses of prostate organoids and tissue-specific mouse models, we investigate whether AR and GR cooperate in the normal prostate, how the two receptors associate and organize within the nucleus, and how their interplay shapes the prostate transcriptional program.

## Results

### GR co-localizes with AR in nuclear foci of prostate epithelial cells and enhances AR phase separation

To characterize the subcellular distribution of AR and GR under physiological conditions, we first examined their expression pattern in the adult mouse prostate. Immunofluorescence analysis of histological sections revealed that both AR and GR are expressed in prostate epithelial cells and localize in nuclear foci (Fig. 1A, B and Fig. S1A). Dual-colour confocal imaging showed extensive overlap between AR- and GR-positive nuclear foci. Quantification using Manders’ overlap coefficients demonstrated that 94.6 ± 0.8% of the GR signal overlapped with AR (M1 = 0.946 ± 0.008), whereas 96.8 ± 0.6% of the AR signal overlapped with GR (M2 = 0.968 ± 0.006) (Fig. 1B), indicating that the two receptors occupy largely shared nuclear compartments in prostate epithelial cells.

**Figure 1.**
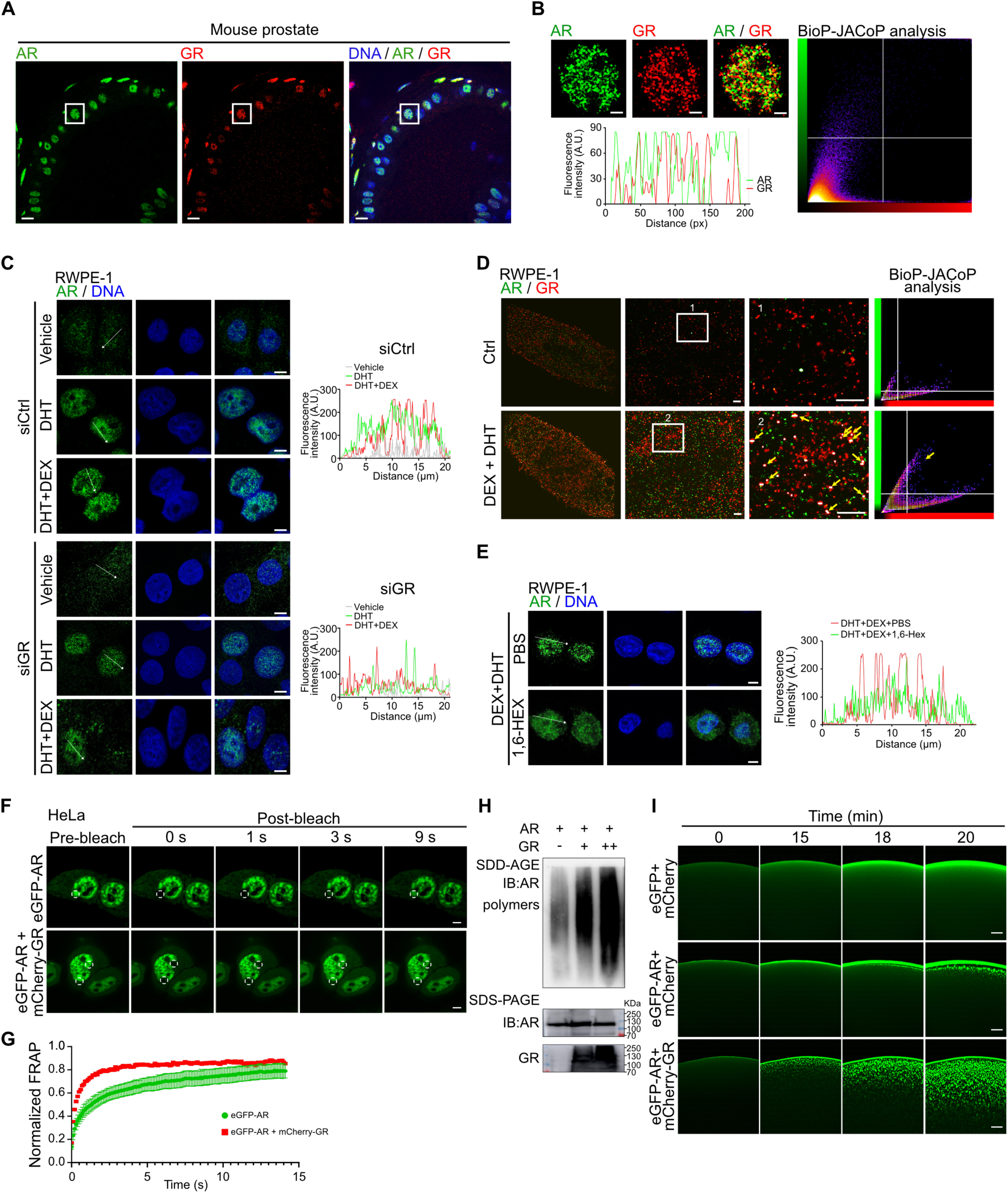
Nuclear co-localization and biophysical properties of AR and GR in prostate. A-B. Confocal images of AR (green) and GR (red) in mouse prostate (A), and per-cell quantification (B). Nuclei are stained with DAPI. Co-localization was scored with BIOP-JACoP (Manders’ coefficients). Scale bars, 50 μm (A), 5 μm (B). C. Immunofluorescence and fluorescence-intensity quantification of AR (green) in RWPE-1 cells treated for 24-h with vehicle (Ctrl), 10 nM DHT or 10 nM DHT + 10 nM DEX, after transfection with GR-targeting siRNA (siGR) or a scramble control siRNA (siCtrl). Nuclei are stained with DAPI. Scale bar, 10 μm. D. Super-resolution SMLM imaging of RWPE-1 cells treated with vehicle, 10 nM DHT and/or 10 nM DEX for 24-h. AR (green) and GR (red). Enlarged views of the boxed regions are shown on the right. Yellow arrows, AR-GR co-localized foci. Co-localization was scored with BIOP-JACoP as in (A, B). Scale bars, 1 µm (overview), 200 nm (insets). E. Immunofluorescence and fluorescence-intensity quantification of AR (green) in RWPE-1 cells treated with 10 nM DHT + 10 nM DEX, with or without 10% 1,6-HEX. Nuclei are stained with DAPI. Scale bar, 10 μm. F-G. FRAP of HeLa cells expressing eGFP-AR and/or mCherry-GR (F), and recovery quantification (G). Scale bar, 5 μm. H. SDD-AGE of recombinant AR with increasing recombinant GR, immunoblotted for AR. Lower panels, SDS-PAGE of input AR and GR. I. Time-lapse images of *in vitro* droplet formation by recombinant eGFP-AR with mCherry or mCherry-GR; eGFP + mCherry, double-negative control. Scale bars, 10 μm.

We next examined the effects of cognate ligand stimulation on AR and GR cellular distribution in RWPE-1 non-tumoral human prostate epithelial cells. While AR was detected in both cytoplasmic and nuclear compartments in the absence of ligand, a 24-h treatment with the AR agonist 5α-dihydrotestosterone (DHT) enhanced its nuclear localization within foci (Fig. 1C), in agreement with previous reports^39^. Combined treatment with DHT and the GR agonist dexamethasone (DEX) further enhanced the number of AR foci, without affecting AR protein levels (Fig. 1C and Fig. S1B). This effect was abolished by GR knockdown (Fig. 1C and S1C), indicating that ligand-activated GR promotes DHT-induced AR nuclear foci formation. Importantly, single-molecule localization microscopy (SMLM) revealed an overlap of AR and GR foci in RWPE-1 cells co-treated with their respective ligands (Fig. 1D), with a median diameter of 80 nm, comparable to that reported for transcriptional condensates containing RNA polymerase II or Mediator complexes^40^. To assess whether these structures exhibit condensate-like properties, cells were treated with 10% 1,6-hexanediol (1,6-HEX), which disrupts weak hydrophobic interactions that contribute to biomolecular condensate formation. Treatment with 10% 1,6-HEX dispersed AR nuclear foci into a diffuse nuclear signal (Fig. 1E), consistent with liquid-like assemblies formed through LLPS.

To assess whether GR influences the properties of AR assemblies, we performed fluorescence recovery after photobleaching (FRAP) in HeLa cells co-expressing eGFP-AR with either mCherry-GR or an mCherry control. AR condensates recovered substantially faster in the presence of GR (t₁_/_₂ = 2.89 s, mobile fraction = 85 %) than in its absence (t₁_/_₂ = 8.35 s, mobile fraction = 73 %) (Fig. 1F and G), indicating that GR promotes AR mobility within the nucleus. Moreover, semi-denaturing detergent agarose gel electrophoresis (SDD-AGE) analysis revealed that increasing GR concentrations enhance AR polymer-like assemblies (Fig. 1H), further supporting a role for GR in promoting AR higher-order oligomerization.

We next investigated the direct influence of GR on AR condensate formation *in vitro*. Recombinant eGFP-AR formed micron-sized spherical droplets *in vitro* within 18 min in the presence of DHT and 10% PEG, and their number and size increased over time (Fig. 1I), showing that *in vitro* liganded AR undergoes LLPS, in agreement with previous reports^31^. By contrast, mCherry-GR formed negligible droplets under identical crowding conditions in the presence of DEX (Fig. S1D), indicating that GR has a low intrinsic propensity to undergo LLPS under these conditions. Co-incubation of eGFP-AR and mCherry-GR in the presence of cognate ligands accelerated AR condensate formation. eGFP-AR-containing droplets were readily detected within 15 min, and were markedly more abundant by 20 min, whereas mCherry-GR-containing droplets were not detected until approximately 40 min (Fig. 1I, S1D and E). To determine whether the GR ligand-binding domain (LBD) is required for this activity, we performed the assay using either full-length mCherry-GR or an LBD deletion mutant (mCherry- GR^ΔLBD^). While full-length GR accelerated AR droplet formation, mCherry-GR^ΔLBD^ failed to enhance eGFP-AR condensate formation (Fig. S1F), indicating that the GR LBD is required for this effect. Together, these findings show that AR and GR co-localize within nuclear foci in prostate epithelial cells and identify a direct role for GR in promoting AR condensate formation. Combined with the FRAP and SDD-AGE analyses, these data further indicate that GR enhances the assembly and molecular dynamics of AR condensates through an LBD-dependent mechanism.

### AR and GR binding sites extensively overlap in mouse prostate

To define the genomic binding landscapes of AR and GR in the mouse prostate, we performed cistrome analyses by chromatin immunoprecipitation followed by sequencing (ChIP-seq). We identified 21,274 AR-binding sites (ARBS) and 21,407 GR-binding sites (GRBS), which were equally distributed across promoter, intronic and intergenic regions (Fig. S2A). More than 90 % of the 11,767 genes bound by AR and the 11,059 genes bound by GR were associated with at least one H3K4me2-marked region, a hallmark of accessible promoter and enhancer regulatory elements, indicating that AR and GR predominantly occupy accessible regulatory elements rather than non-functional genomic sites in the mouse prostate (Fig. S2B). Chromatin-state annotation showed that approximately one-third of ARBS and one-fifth of GRBS were located to active promoters, marked by high H3K4me3/2 and low H3K4me1 signals, whereas the remaining peaks mapped predominantly to active enhancers, enriched for H3K4me1/2 and depleted of H3K4me3 histone marks (Fig. 2A and B). *De novo* motif analysis revealed a strong enrichment of the canonical HRE at enhancer-associated ARBS and GRBS, whereas promoter-proximal binding sites encompassed motifs for promoter-associated transcription factors, including NFY (Fig. 2A and B).

**Figure 2.**
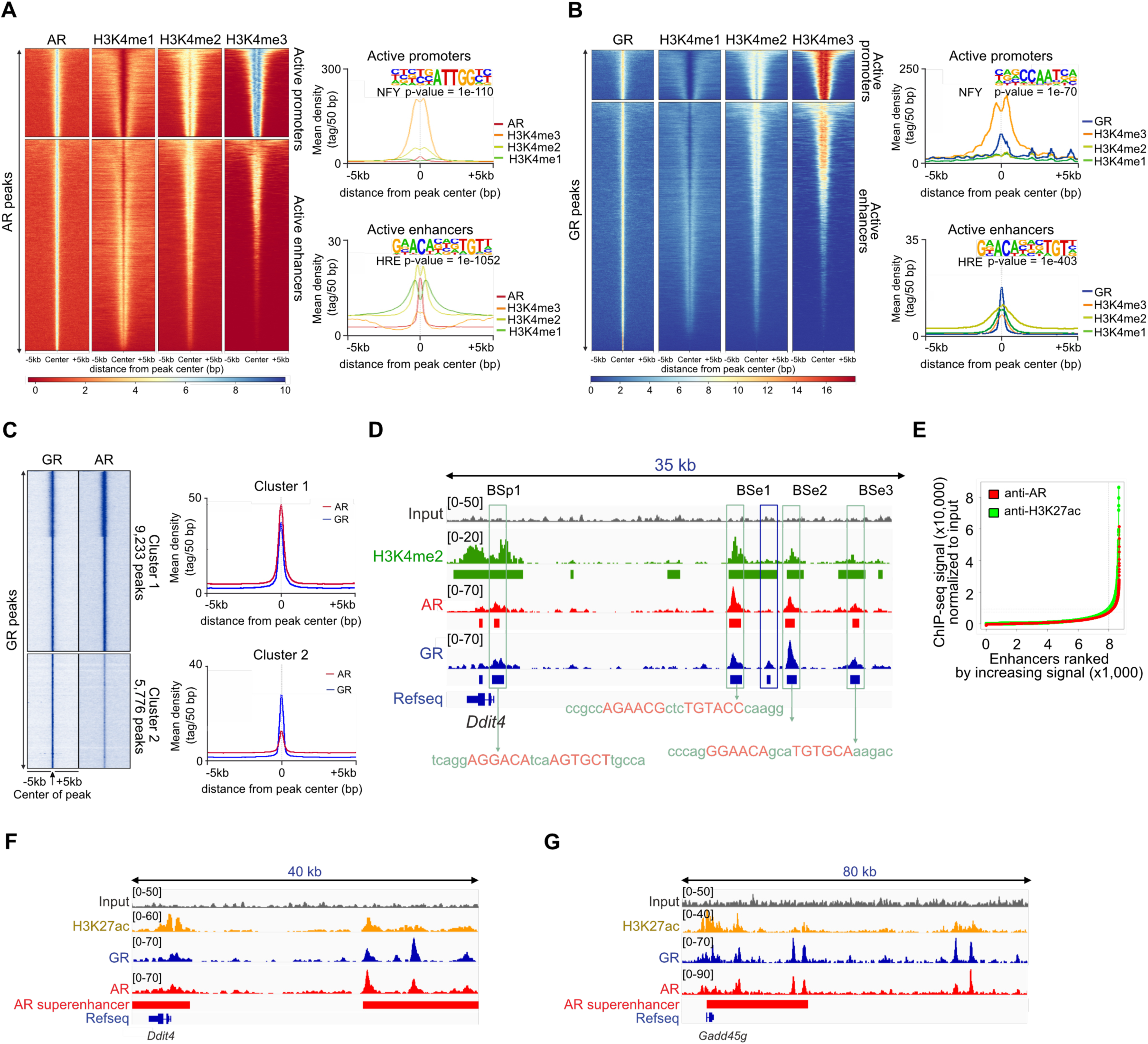
Genome-wide occupancy of AR and GR in mouse prostate. A-B. Tag density heatmaps of AR (A) and GR (B) ChIP-seq peaks with H3K4me1, H3K4me2 and H3K4me3 within ± 5 kb of peak centre, with average tag-density profiles of active promoters and active enhancers and corresponding *de novo* motif analysis. C. Heatmap of AR and GR ChIP-seq signal centred on GR peaks, k-means clustered into Cluster 1 (9,233 peaks, AR-GR co-occupancy) and Cluster 2 (5,776 peaks, GR alone). Average tag density profiles at right. The two first clusters were considered as Cluster 1. D. AR, GR and H3K4me2 occupancy at the *Ddit4* locus. Enhancer elements (BSe1-BSe3) and the promoter region (BSp1) are indicated in green boxes marking the HRE-containing sequence. E. AR and H3K27ac ChIP-seq signal (reads per million, RPM) across all mouse-prostate enhancers, ranked by increasing AR signal (super-enhancer ranking). F-G. IGV browser tracks of H3K27ac, GR, AR ChIP-seq signal and the annotated AR super-enhancer at the *Ddit4* (F) or *Gadd45g* (G) loci.

Notably, almost 60 % of AR-binding sites overlapped with those of GR (Fig. S2C), and approximately 70 % of AR and GR target genes were common (Fig. S2D). SeqMINER analysis further showed that GR signal was enriched at two-thirds of AR peaks (Fig. S2E, cluster 1), and about 60 % of GR binding sites exhibited coincident AR enrichment (Fig. 2C). Consistent with this extensive co-occupancy, shared AR-GR binding sites displayed higher average ChIP-seq signal than receptor-specific sites (Fig. 2D). Moreover, approximately 60 % of AR- or GR-specific peaks were located within 50 kb of a shared AR-GR binding site (Fig. 2D, S2F and Table S1), indicating that AR and GR occupy a largely overlapping cis-regulatory landscape in the mouse prostate, with receptor-specific binding events frequently occurring in the vicinity of shared regulatory elements.

To investigate whether AR and GR co-occupy extended enhancer domains, we identified super-enhancers using the ROSE algorithm^41,42^, based on H3K27ac enrichment, with AR serving as the lineage-defining transcription factor for prostate epithelial cells^36,43–45^. This analysis identified 245 AR-associated super-enhancers, all of which were also bound by GR (Fig. 2E, S2G and H and Table S2), as exemplified for *Ddit4* and *Gadd45g* (Fig. 2F and G). Altogether, these findings demonstrate that AR and GR are co-recruited to cis-regulatory elements encompassing HREs in mouse prostate, including active enhancers and AR-associated super-enhancers.

### AR and GR are embedded in similar coactivator and corepressor complexes at chromatin

To characterize the AR- and GR-containing chromatin-associated macromolecular complexes, we performed rapid immunoprecipitation mass spectrometry (RIME)^46^ with anti-AR or anti-GR antibodies on nuclear extracts from RWPE-1 cells co-treated with DEX and DHT. GR-RIME identified 174 significantly enriched proteins, with GR (NR3C1) recovered as the top-ranked hit, thereby validating the specificity of our approach (Fig. 3A and Table S3). STRING network analysis revealed interactions with established nuclear receptor coactivators, including EP300, NCOA2/TIF2, NCOA3/SRC3, KDM1A/LSD1 and STAT3, as well as epithelial transcription factors such as GRHL2, BNC1 and TP63 (Fig. 3A). The interaction between GR and TP63 was independently confirmed by co-immunoprecipitation experiments on prostate nuclear extracts (Fig. S3A). In parallel, AR-RIME identified 143 significantly enriched proteins (Fig. 3B and Table S4), amongst which the AP-1/bZIP factors FOSL2, JUNB, FOS, ATF3 and CEBPB, the ETS factor ERF, and components of the RNA polymerase II machinery (e.g. POLR2C) (Fig. S3B).

**Figure 3.**
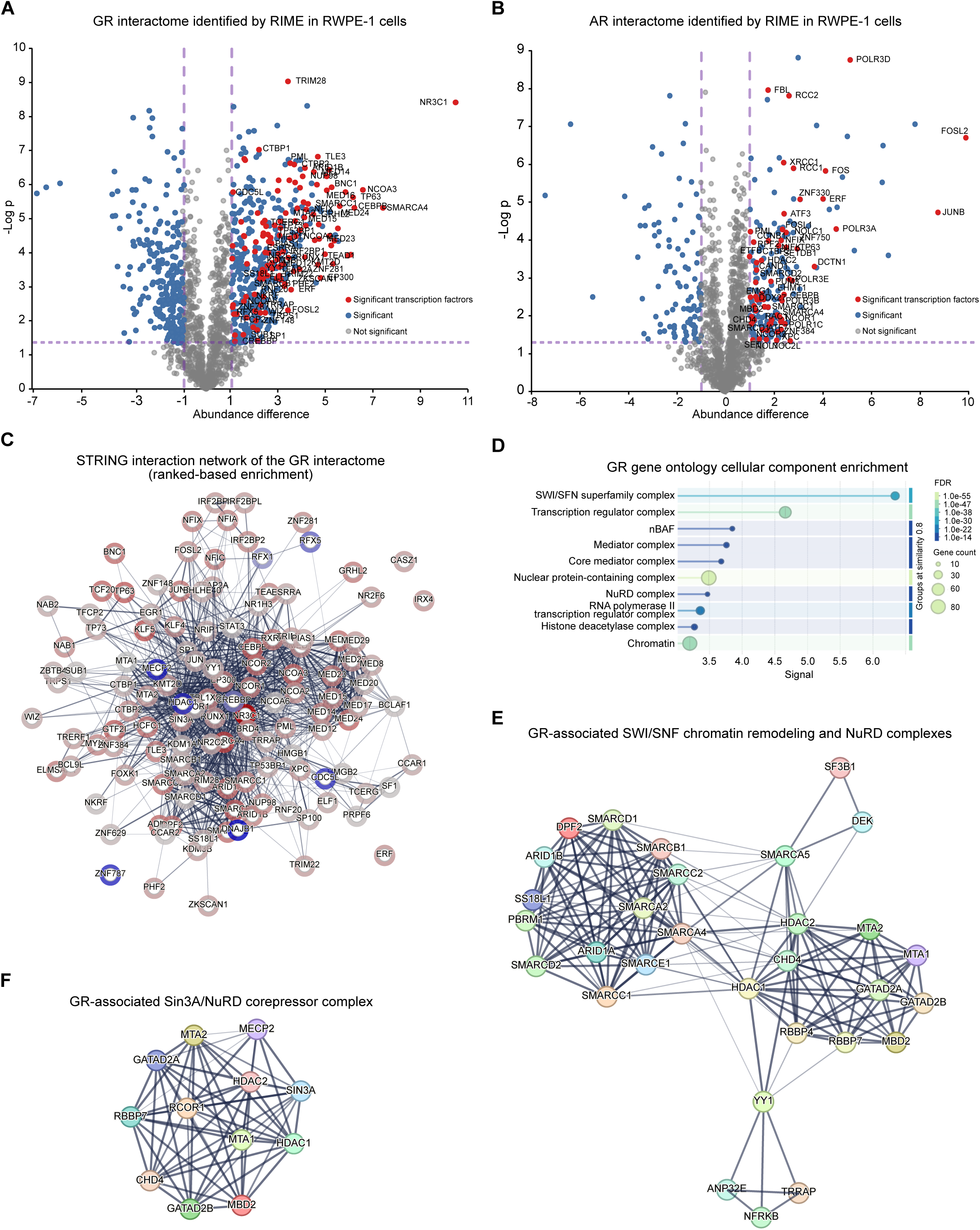
The AR and GR interactomes and their associated chromatin coregulator complexes. A-B. Volcano plot of protein enrichment in GR-RIME (A) and AR-RIME (B) from RWPE-1 cells co-treated with DHT and DEX, relative to IgG control (n = 3 biological replicates per condition). The x-axis shows the log₂ enrichment (RIME/IgG) and the y-axis the -log_10_ P value (two-sided Student’s t-test). Proteins passing the significance threshold (permutation-based FDR<0.05, Perseus) are coloured: transcription factors, red; other significant interactors, blue; non-significant proteins, grey. C. STRING protein-protein interaction network of significant GR-enriched proteins together with their STRING-predicted partners (confidence score≥0.4, 454 nodes). Red node outlines, transcription factors. Blue node outlines, chromatin- and transcription-regulator complex components. D. Gene Ontology Cellular Component enrichment of the GR interactome. Dot size represents number of proteins per term and colour encodes FDR. E-F. STRING subnetworks of GR-associated SWI/SNF (BAF/nBAF) chromatin remodelling and NuRD complexes (E), and of the SIN3A and NuRD corepressor complexes (F).

Rank-based STRING network analysis of the GR interactome yielded a highly interconnected network comprising 454 proteins (Fig. 3C). Gene Ontology enrichment analysis revealed a significant association with chromatin regulatory complexes, including the SWI/SNF (BAF/nBAF), the Mediator complex, and the RNA polymerase II transcription regulator complexes, as well as the NuRD and histone deacetylase repressive complexes (Fig. 3D). Consistent with these enrichments, GR connected to core components of the SWI/SNF complex (e.g. SMARCA4, SMARCC1/2, ARID1A/B), the Mediator complex (e.g. MED1, MED8, MED12, Fig. S3D), and the SIN3/NuRD/NCoR repressive machinery (e.g. HDAC1/2, CHD4, MTA1/2, RBBP4/7, SIN3A, MECP2, NCOR1, GATAD2A/B, MBD2, DEK) (Fig. 3E, F and S3C, D). The AR interactome exhibited a highly similar organization, encompassing SWI/SNF subnetwork (e.g. SMARCA4, SMARCC1, SMARCD1/2, PBRM1, DPF2), together with NCoR/NuRD-associated corepressors (e.g. NCOR1/2, HDAC2, CHD4, MBD2, CTBP2, TLE3 and TRIM28) (Fig. S3F and G). Thus, despite distinct receptor-specific interactors, both AR and GR associate with a common set of chromatin remodelling, transcriptional coactivator, and corepressor complexes, indicating that the two receptors are embedded within a shared chromatin regulatory environment.

### AR and GR form heterodimers through the GR ligand-binding domain

Neither RIME dataset recovered the reciprocal receptor (AR was not detected from GR-RIME and GR from AR-RIME). Because transient or sub-stoichiometric chromatin-associated interactions are often underrepresented in unbiased affinity purification-mass spectrometry experiments^47^, we tested whether AR and GR directly interact using targeted biochemical approaches. Co-immunoprecipitation from mouse prostate and RWPE-1 nuclear extracts confirmed an association between endogenous AR and GR (Fig. 4A and S4A). To determine whether this interaction is direct, we performed *in vitro* pull-down assays, using recombinant MBP-tagged GR and Avi-tagged AR. MBP-GR efficiently retained AR, demonstrating a direct interaction between the two receptors (Fig. 4B). To map the specific GR domains mediating this interaction, we co-transfected Flag-tagged full-length or truncated GR variants (Fig. 4C) with Myc-tagged AR in HEK293T cells in the presence of ligand. Immunoprecipitation experiments showed that recombinant AR and GR interact, and identified the GR ligand-binding domain (GR^LBD^) as necessary for AR association (Fig. 4D). Consistent with this result, GST pull-down assays revealed that the GR^LBD^ was sufficient to bind AR *in vitro* (Fig. 4E and F), showing that AR directly associates with GR *via* the GR^LBD^.

**Figure 4.**
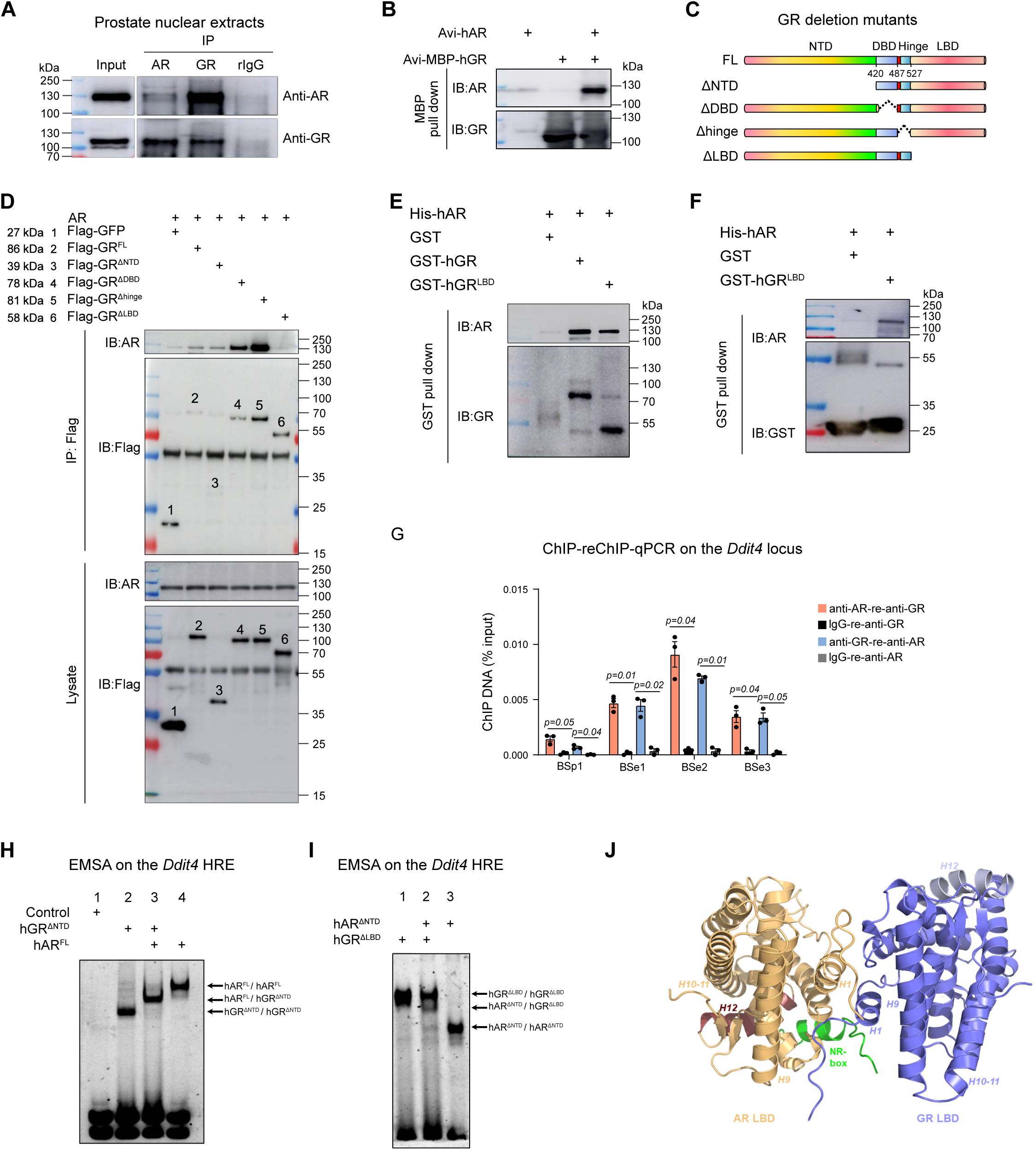
Physical interaction of AR and GR through the GR ligand-binding domain. A. Co-immunoprecipitation of AR and GR in mouse prostate nuclear lysates with anti-AR or anti-GR antibodies, immunoblotted for both. Rabbit IgG (rIgG) served as a control. 10% input were also analysed. Different protein amounts were loaded per lane to avoid overexposure. B. *In vitro* amylose MBP pull-down of Avi-MBP-GR and Avi-AR co-expressed in Sf21 cells in the presence of DEX and DHT, immunoblotted for AR and GR. AR expressed alone, control for non-specific binding to amylose resin. C. Schematic domain structure of human GR full-length (FL) and deletion mutants. Intrinsically disordered NTD, DBD, hinge and LBD, with residue boundaries indicated. ΔNTD, ΔDBD, Δhinge, and ΔLBD lack the corresponding domains. D. Anti-Flag immunoprecipitation from HEK293T cells co-expressing full-length AR with full-length or deletion-mutant Flag-GR, immunoblotted for AR (top) and Flag (bottom). Total lysate was used as control for vector expression. E-F. GST pull-down of purified His-hAR with GST-hGR (FL), hGR-LBD or GST alone (all from *E. coli*), immunoblotted for AR and GR (E) or AR and GST (F). G. Sequential ChIP (ChIP-reChIP-qPCR) on wild-type prostate nuclear extracts at the *Ddit4* GR-binding sites with the indicated antibodies. Mean ± SEM, two-tailed Mann-Whitney test. H. EMSA on the *Ddit4* BSe2 HRE (5’-AGAACAttgTGTTCT-3’) with whole-cell extracts from HEK293T cells expressing Flag-hGR^ΔNTD^ and/or hAR^FL^ in the presence of ligands. Lane 1, mock-transfected. Lane 2, hGR^ΔNTD^. Lane 3, hGR^ΔNTD^ + hAR^FL^. Lane 4, hAR^FL^. Homodimeric (hGR^ΔNTD^/hGR^ΔNTD^, hAR^FL^/hAR^FL^) and heterodimeric (hAR^FL^/hGR^ΔNTD^) protein-DNA complexes indicated at right. I. EMSA as in (H) with extracts expressing Myc-hAR^ΔNTD^ and/or Flag-hGR^ΔLBD^. Lane 1, hGR^ΔLBD^. Lane 2, hAR^ΔNTD^ + hGR^ΔLBD^. Lane 3, hAR^ΔNTD^. The heterodimer band in lane 2 is reduced relative to the corresponding in (H, lane 3), whereas both truncated receptors form homodimers. J. AlphaFold 3 model of an AR-LBD (yellow)-GR-LBD (blue) heterodimer in the presence of an NR-box (LxxLL) peptide (green).

To determine whether AR and GR co-occupy the same regulatory regions, we considered the *Ddit4* locus, which contains four shared AR- and GR-binding sites encompassing HREs, separated by at least 5 kb (Fig. 2D). ChIP-qPCR confirmed AR and GR occupancy at these regions in the mouse prostate (Fig. S4B). Importantly, ChIP-reChIP assays revealed that AR and GR are co-recruited to all four sites of the *Ddit4* locus under physiological conditions (Fig. 4G).

To decipher whether AR and GR can form DNA-bound complexes *in vitro*, we performed electrophoretic mobility shift assays (EMSA) using whole-cell extracts from HEK293T cells expressing AR and GR variants, together with an oligonucleotide probe spanning the *Ddit4* HRE (5’-AGAACAttgTGTTCT-3’)^48^, in the presence of their respective ligands. Since the DNA complexes formed by full-length AR and GR (hAR^FL^ and hGR^FL^) exhibited similar electrophoretic mobility (Fig. S4C, lanes 2-3), we used an N-terminally truncated GR variant (hGR^ΔNTD^), which forms a faster-migrating complex, thereby improving resolution of distinct species (Fig. 4H, lane 2, and Fig. S4C, lane 6). Co-incubation of hAR^FL^ and hGR^ΔNTD^ predominantly yielded a protein-DNA complex of intermediate position relative to the respective homodimeric complexes (Fig. 4H, lanes 2, 3 and 4), consistent with formation of a heterodimeric protein-DNA bound complex. The reciprocal EMSA experiment using AR^ΔNTD^ and hGR^FL^ resulted in a similar intermediate-mobility species (Fig. S4C). In contrast, heterodimer formation was markedly reduced when hAR^ΔNTD^ was co-incubated with GR^ΔLBD^ (Fig. 4I, lane 2), despite robust homodimer formation by each receptor on the probe (Fig. 4I, lanes 1 and 3, respectively), further indicating that the GR ligand-binding domain contributes to AR-GR complex formation on DNA. A similar outcome was obtained using the *Tmprss2* HRE (5’-GGAACTcttTGTTCA-3’) as a probe (Fig. S4D), indicating that AR-GR heterodimer formation is not restricted to the *Ddit4* element. Together, these data show that GR forms heterodimeric complexes with AR on the DNA in a GR-LBD-dependent manner.

To gain insights into the structural basis of the AR-GR heterodimer, we modelled the arrangement of their LBDs in complex with a short LxxLL motif-containing peptide using AlphaFold 3 to stabilize the interacting surfaces between the two LBD. The resulting predictions consistently identified a heterodimeric interface primarily mediated by helices H1 and H3 of both LBD subunits, with robust confidence metrics: a predicted template modelling (pTM) score of 0.54 and an interface pTM (ipTM) score of 0.73 (Fig. 4J). Notably, this arrangement resembles the dimerization interface observed in the crystal structure of the multidomain GR(385-777)-DNA complex^49,50^, indicating that heterodimer formation may exploit an interface geometry that is related to, yet distinct from, AR and GR homodimer configurations. Together, these data support a model in which AR and GR form ligand-dependent heterodimers on DNA through direct interactions mediated primarily by their ligand-binding domains.

### AR-GR complexes exert DNA-dependent transcriptional regulation

To investigate the AR and GR transcriptional activities in a physiologically relevant model, we analysed mouse prostate epithelial organoids, cultured in androgen-containing medium to ensure their survival and growth. Immunofluorescent detection of AR and GR revealed that the two receptors co-localize within nuclear foci (Fig. 5A). Consistent with our observations made in RWPE-1 cells, the number and intensity of AR and GR foci increased upon DEX treatment, whereas exposure to 10% 1,6-HEX dispersed these structures (Fig. 5A, B and S5A). These findings indicate that AR-GR nuclear assemblies formed in prostate organoids exhibit condensate-like properties.

**Figure 5.**
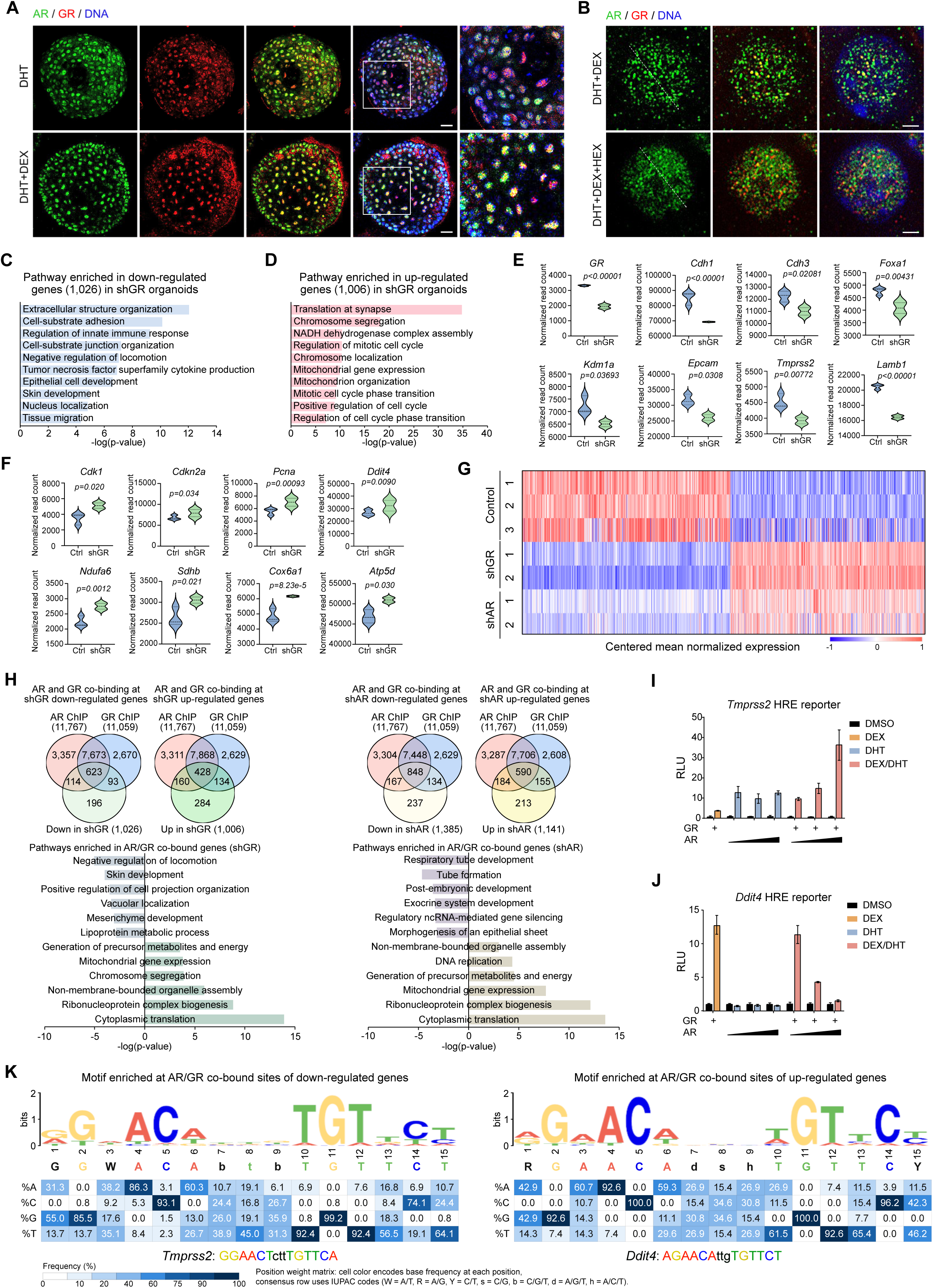
AR- and GR-dependent transcriptional programs and the response elements underlying them in mouse prostate organoids. A. Immunofluorescence of AR (green) and GR (red) in wild-type mouse prostate organoids treated with DHT or DHT+DEX. Nuclei are stained with DAPI. Scale bars, 50 μm; insets, 10 μm. B. Immunofluorescence of AR (green) and GR (red) in organoids treated with DHT + DEX in the presence or absence of 10% 1,6-HEX. Nuclei are stained with DAPI. Scale bar, 50 μm. C-D. Pathway enrichment for genes down- (C) and up-regulated (D) in organoids transduced with GR-targeting shRNA versus control shRNA. P-values are adjusted with the Benjamini-Hochberg method. E-F. Normalized RNA-seq read count (violin plots) of representative down- (E) or up-regulated (F) genes after GR knockdown. *P*-values were obtained from DESeq2. G. Heatmap of mean-centred normalized expression of down- and up-regulated genes in organoids transduced with shRNA against GR, AR or control. H. Venn diagrams of the overlap between AR- or GR-bound genes (ChIP-seq) and genes differentially expressed after GR or AR knockdown, with KEGG enrichment of the co-bound genes. I-J. Transactivation assay in HEK293T cells transfected with constant hGR and increasing hAR plus a luciferase reporter driven by the *Tmprss2*-type (I) or *Ddit4*-type (J) HRE, treated with vehicle (DMSO), 100 nM DEX, 100 nM DHT or DEX+DHT. Luciferase activity normalized to eGFP and to the vehicle control. K. *De novo* motif discovery at AR/GR ChIP-seq peaks linked to genes differentially expressed after AR or GR silencing. Left, activating *Tmprss2*-type consensus (5’-GGWACAbtbTGTTCT-3’) at co-bound sites of genes down-regulated upon depletion (e.g. *Tmprss2*). Right, repressive *Ddit4*-type consensus (5’-RGAACAdshTGTTCY-3’) at co-bound sites of genes up-regulated upon depletion (e.g. *Ddit4*). Top row, sequence logo (bits). Lower rows, position-specific base-frequency matrix (% A/C/G/T, colour-coded). Consensus in IUPAC code (W=A/T, R=A/G, Y=C/T, S=C/G, b=C/G/T, d=A/G/T, h=A/C/T).

We next examined the AR and GR transcriptomes in primary prostate organoids following lentiviral shRNA-mediated knockdown of either receptor (Fig. S5B and C). GR silencing resulted in 1,026 down-regulated genes (reads >50, p-value<0.05), mainly involved in cell motility and epithelium development, including *Epcam*, cadherins *Cdh1* and *Cdh3*, laminins (e.g. *Lama3* and *Lamb1*), as well as *Tmprss2*, *Nkx3-1*, *Foxa1* and *Kdm1a* (also known as *Lsd1*). We also identified 1,006 up-regulated genes upon GR knockdown implicated in cell cycle (e.g. *Cdk1*, *Cdkn2a*, *Pcna*, and centromere proteins of the CENP family) and in oxidative phosphorylation, including nuclear-encoded subunits of the mitochondrial respiratory complexes (e.g. *Ndufa6, Sdhb, Uqcrb, Cox6a1,* and *Atp5d*) (Fig. 5C-F). *Ddit4* was likewise up-regulated in the organoid RNA-seq dataset (Fig. 5F).

AR silencing resulted in the down-regulation of 1,385 genes and up-regulation of 1,141 genes, with pathway enrichment profiles closely mirroring those observed upon GR knockdown (Fig. 5G). This transcriptomic convergence between AR- and GR-dependent transcriptional programs indicates that the two receptors regulate a substantial fraction of common gene networks in prostate epithelial cells. RT-qPCR independently validated RNA-seq data on representative target genes, including reduced *Tmprss2* and increased *Ddit4* expression following depletion of either receptor (Fig. S5D).

Integration of the cistrome and transcriptome datasets revealed that most of the genes differentially expressed upon AR or GR silencing are bound by both receptors. Among the genes down-regulated upon GR knockdown, ∼60 % were bound by both AR and GR, while only ∼10 % were associated with AR- or GR-specific binding events (Fig. 5H). Similar observations were made for genes up-regulated following GR knockdown, as well as for genes differentially expressed upon AR silencing, indicating that the transcriptional outputs are predominantly associated with the shared AR-GR cistrome rather than receptor-specific occupancy, as exemplified for *Ddit4*, *Gadd45g*, *Nkx3-1*, and *Tmprss2* (Fig. 5H). Pathway analysis on the shared target genes showed that genes down-regulated following receptor silencing were enriched for epithelial development and cell adhesion programs, whereas up-regulated targets converged on cytoplasmic translation, mitochondrial gene expression, and DNA replication pathways (Fig. 5H). Together, these findings indicate the shared AR-GR cistrome translates into transcriptional programs that promote epithelial identity while restraining proliferative and metabolic pathways.

To determine whether AR and GR exert context-dependent transcriptional responses at shared DNA response elements, we conducted transactivation assays in HEK293T cells co-expressing hAR and/or hGR together with HRE-driven reporters in the presence of cognate ligands. Reporters harbouring two tandem copies of an HRE from either the *Tmprss2* locus (5ʹ-GGAACTcttTGTTCA-3ʹ, historically annotated as a canonical ARE) or the *Ddit4* locus (5ʹ-AGAACAttgTGTTCT-3ʹ, historically annotated as a canonical GRE)^48^, were positioned upstream of a minimal promoter. At the *Tmprss2* HRE-containing reporter, hAR expression led to the induction of luciferase activity in response to DHT, which was further enhanced upon hGR co-expression in the presence of DHT and DEX (Fig. 5I). In sharp contrast, the *Ddit4* HRE-driven reporter was robustly activated by GR following DEX treatment, whereas liganded hAR had little effect. Notably, co-expression of AR progressively attenuated GR-dependent activation in the presence of DHT in a dose-dependent manner (Fig. 5J). Together, these results indicate that the transcriptional output of AR-GR complexes is determined by the underlying HRE sequence. Whereas AR and GR cooperate to enhance transcription from the *Tmprss2* HRE, AR antagonizes GR-mediated activation at the *Ddit4* HRE, indicating that closely related HREs can encode distinct transcriptional responses to AR-GR complexes.

To assess whether the sequence-dependent effects of AR-GR complexes extend genome-wide, we intersected the genes differentially expressed upon AR or GR silencing with the genes associated with shared AR-GR binding sites, and performed MEME-ChIP motif enrichment on the corresponding co-bound peak regions. Peaks associated with genes down-regulated upon silencing (i.e. activated by the AR-GR complex) were significantly enriched for the element 5ʹ-GGWACAbtbTGTTCT-3ʹ, which closely matches the *Tmprss2* HRE. In contrast, peaks linked to genes up-regulated upon receptor knockdown (i.e. repressed by the AR-GR complex) harboured the 5ʹ-RGAACAdshTGTTCY-3ʹ motif, corresponding to the *Ddit4* HRE (Fig. 5K). Together with the reporter assays, these data support a model in which subtle sequence variation within HREs specifies the transcriptional output of AR-GR complexes, distinguishing the loci they activate from those they repress.

### Epithelial GR ablation derepresses metabolic programs and impairs prostate homeostasis

To determine the physiological role of GR in prostate epithelial cells *in vivo,* we generated GR^(i)pe-/-^mice, in which GR is selectively deleted in prostate luminal cells after puberty, by crossing PSA-CreER^T2^ mice^51^ with GR^L2/L2^ (floxed *Nr3c1*) mice^48^ and inducing recombination by tamoxifen administration at 8 weeks of age. Tamoxifen-treated littermates lacking the PSA-CreER^T2^ transgene served as controls (Fig. S6A and B). Immunofluorescence staining confirmed the loss of GR specifically in luminal cells of the dorsolateral prostate (DLP) of GR^(i)pe-/-^ mice, while AR expression persisted (Fig. 6A and S6C). GR-deficient luminal cells exhibited a more diffuse AR nuclear distribution, whereas AR remained located in nuclear foci in basal cells lacking the CreER^T2^ recombinase (Fig. 6B and S6D). This phenotype recapitulated the GR-dependent enhancement of AR nuclear foci formation observed in RWPE-1 cells (Fig. 1C), further supporting that GR promotes AR nuclear assembly *in vivo*.

**Figure 6.**
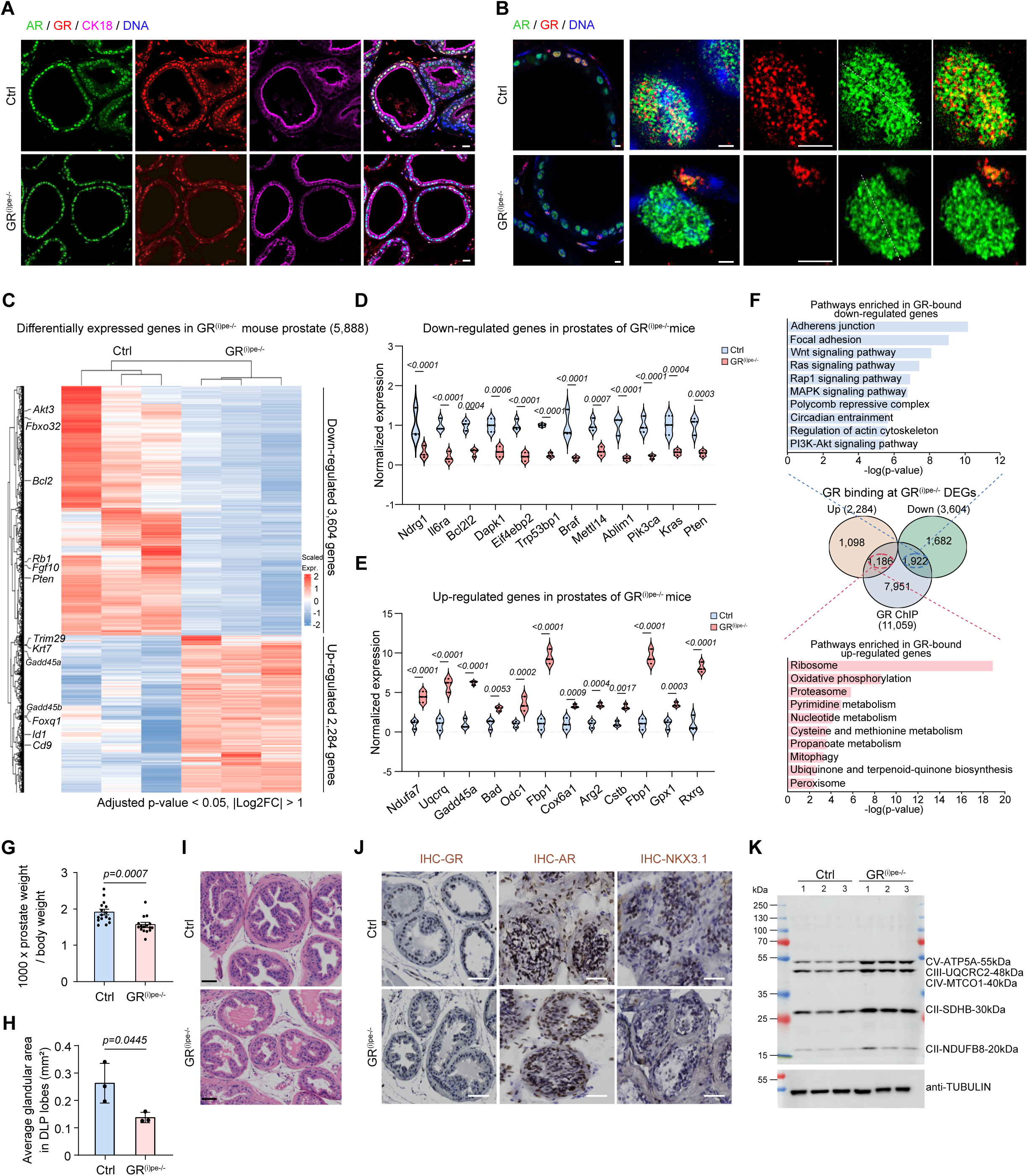
Consequences of prostate-epithelium-specific GR ablation for AR-dependent transcription and tissue homeostasis *in vivo*. A. Immunofluorescence of AR (green), GR (red) and CK18 (magenta) in control and GR^(i)pe-/-^ mouse prostate. Nuclei are stained with DAPI. Scale bars, 50 μm. B. High-magnification single-cell nuclear distribution of AR (green) and GR (red) in control and GR^(i)pe-/-^prostate. Nuclei are stained with DAPI. Scale bar, 5 μm. C. Heatmap of scaled expression of differentially expressed genes in control versus GR^(i)pe-/-^ prostates. Selected drivers annotated at left. D-E. Normalized expression (violin plots) of representative down- (D) or up-regulated (E) genes in GR^(i)pe-/-^ prostate. *P*-values were obtained from DESeq2. F. Venn diagram of the overlap between up- and down-regulated genes in GR^(i)pe-/-^ prostates and GR ChIP-seq target genes, with KEGG enrichment of up-regulated (left, red) and down-regulated (right, blue) GR-bound genes. G. Relative prostate weight (prostate weight × 1,000 / body weight) in control and GR^(i)pe-/-^ mice. Mean ± SEM, two-tailed Mann-Whitney test. n = 16 mice per group. H. Mean glandular area (mm²) in the dorsolateral prostate (DLP) of control and GR^(i)pe-/-^ mice. Mean ± SEM, two-tailed Mann-Whitney test, n = 3 mice per group. I. Representative H&E-stained prostate sections from control and GR^(i)pe-/-^ mice. Scale bar, 50 μm. J. Immunohistochemistry for GR, AR and NKX3.1 in control (top) and GR^(i)pe-/-^ (bottom) prostate. Scale bar, 50 μm. K. Representative immunoblot of the indicated OXPHOS-complex subunits in prostates of control and GR^(i)pe-/-^ mice. α-Tubulin was used as a loading control. n = 3 mice per group.

To characterize the transcriptional consequences of GR loss in the prostate luminal cells, we performed RNA-seq on control and GR^(i)pe-/-^ prostates. This analysis identified 5,888 significantly differentially expressed genes (p-adj <0.05, reads > 50, |log2FC| >1), comprising 3,604 down-regulated and 2,284 up-regulated transcripts (Fig. 6C). Among down-regulated genes, we identified regulators of epithelial identity and tumour suppression, including *Pten*, *Rb1*, *Bcl2* and *Fgf10*, whereas up-regulated genes encompassed mitochondrial subunits together with metabolic and stress-response factors such as *Ddit4*, *Gadd45a*, *Krt7* and *Trim29* (Fig. 6C and S6E). RT-qPCR on independent samples validated the up-regulation of nuclear-encoded mitochondrial and metabolic genes (*Ndufa7*, *Uqcrq*, *Cox6a1*, *Gpx1*) and the reduction of signalling and survival factors (*Ndrg1*, *Braf*, *Pik3ca*, *Kras*, *Pten*) (Fig. 6D and E), consistent with the transcriptional changes observed following GR silencing in prostate organoids. To assess whether these transcriptional changes following GR loss reflect direct GR activity, we integrated GR cistrome and transcriptome. Of the 2,284 up-regulated genes upon GR deletion, 1,186 (52%) were directly bound by GR, while 1,922 of 3,604 down-regulated genes (53%) harboured GR binding sites (Fig. 6F), indicating that a substantial fraction of GR-dependent transcriptional programs is directly linked to receptor occupancy.

KEGG pathway analysis of the GR-bound genes induced following GR deletion revealed enrichment in ribosome, oxidative phosphorylation, proteasome, pyrimidine and nucleotide metabolism, cysteine and methionine metabolism, propanoate metabolism, peroxisome, mitophagy, and ubiquinone biosynthesis (Fig. 6F), indicating that GR restrains a broad metabolic transcriptional program in prostate epithelial cells under physiological conditions. Conversely, the GR-bound genes reduced following GR deletion were enriched in adherens junction, focal adhesion, Wnt, Ras, MAPK, Rap1, PI3K-Akt and calcium signalling, polycomb repressive complex, circadian entrainment and regulation of the actin cytoskeleton (Fig. 6F), consistent with a role for GR in maintaining epithelial cell signalling programs. To evaluate the convergence of GR-dependent transcriptional programs across our *in vitro* and *in vivo* experimental models, we intersected the transcriptomes of GR-silenced organoids and GR^(i)pe-/-^prostates. This analysis identified 391 commonly up-regulated genes and 425 commonly down-regulated genes between the two systems (Fig. S6F). Gene Ontology analysis of the shared up-regulated genes revealed a predominant enrichment in metabolic and translational programs, including cytoplasmic translation, NADH dehydrogenase complex assembly and mitochondrial gene expression (Fig. S6F), indicating that these programs are consistently de-repressed upon GR loss in both systems. In addition, the shared down-regulated genes were enriched for epithelial structural and signalling programs, including cell-substrate junction organization, epithelial cell development, morphogenesis of an epithelial sheet and actin filament organization (Fig. S6F). These cross-system analyses identify a conserved GR-dependent transcriptional program in prostate epithelial cells, activating epithelial maintenance and adhesion genes while repressing metabolic and translational programs.

Because GR loss altered AR nuclear distribution without abolishing AR expression (Fig. 6B and S6C, D), we investigated whether the dysregulated loci were also occupied by AR. Intersection of the GR^(i)pe-/-^ transcriptome with the AR cistrome showed that AR was bound to a fraction of the differentially expressed genes comparable to that of GR, with 1,220 of the 2,284 up-regulated genes (53%) and 2,017 of the 3,604 down-regulated genes (56%) overlapping AR binding sites (Fig. S6G). KEGG analysis of AR-bound genes revealed the same functional categories as the GR-bound set, including metabolic and translational programs for the up-regulated genes following GR loss and epithelial adhesion and signalling programs for the down-regulated genes (Fig. S6G). Notably, a substantial proportion of the differentially expressed genes was bound by the two receptors (923 up-regulated and 1,570 down-regulated genes, Fig. S6H). The co-bound activated genes were enriched in ribosome, oxidative phosphorylation, thermogenesis, nucleotide and pyrimidine metabolism, proteasome and peroxisome pathways, whereas the co-bound repressed genes were enriched in adherens junction, Wnt, Rap1, MAPK and Ras signalling, focal adhesion, Polycomb repressive complex and prostate cancer pathways (Fig. S6H). Together, these analyses indicate that AR and GR share genomic regions related to metabolic and epithelial transcription programs, differentially expressed upon GR loss in prostate epithelial cells. We next examined the physiological consequences of GR ablation in prostate epithelial cells. One week after GR deletion, the prostate weight relative to body weight and average glandular area in DLP lobes were significantly reduced in GR^(i)pe-/-^ mice compared with controls (Fig. 6G, H, I and S6I). Immunohistochemistry confirmed the loss of GR expression in luminal cells, without affecting AR expression, and revealed a marked reduction in the staining of the AR target NKX3.1 (Fig. 6J). Consistent with the transcriptomic changes, western blot analysis of prostate extracts from GR^(i)pe-/-^ mice revealed elevated levels of mitochondrial respiratory complexes I to V subunits (Fig. 6K). In agreement, Seahorse metabolic flux analysis on RWPE-1 cells silenced for AR or GR revealed an increased basal and maximal mitochondrial respiration and enhanced glycolytic activity relative to control cells (Fig. S6J), providing evidence that receptor loss drives metabolic reprogramming.

Collectively, these data demonstrate that GR in prostate epithelial cells maintains prostate homeostasis by sustaining epithelial identity, while restraining metabolic and cell cycle transcriptional programs.

## Discussion

Since the 1990s, when AR and GR heterodimerization was first reported^24^, numerous studies have sought to dissect how the two receptors act. Our findings reveal an essential and previously underestimated cooperation between AR and GR in the physiological regulation of transcription in prostate epithelium. While AR has long been regarded as the principal driver of prostate epithelial identity^36,43–45,52^, studies in prostate cancer have shown that GR can substitute for AR during androgen-deprivation or AR-targeted therapy, sustaining expression of AR-regulated genes and contributing to anti-androgen resistance^1^. These studies have typically interpreted GR up-regulation as an adaptive response to treatment. Our work instead demonstrates that AR-GR cooperativity is intrinsic to normal prostate biology, establishing GR as a physiological partner of AR rather than merely a pathological surrogate.

Beyond chromatin binding, GR enhances AR nuclear condensation through liquid-liquid phase separation (LLPS), a mechanism increasingly implicated in transcriptional control, particularly for nuclear receptors and for coactivator assemblies involving Mediator and RNA polymerase II^30,40^. Whereas previous studies established that AR and GR can undergo phase separation independently^29,31,34,53^, our *in vitro* reconstitution assays reveal an asymmetric relationship: GR condenses poorly on its own yet accelerates AR droplet formation, increases droplet growth, and enhances AR mobility within condensates in a GR-LBD-dependent manner. Inter-receptor interactions can thus modulate the biophysical properties of transcription factor condensates. Moreover, our proteomic data indicate that the resulting complexes are not exclusively activation hubs, as typically described for Mediator/Pol II condensates. Network and Gene Ontology analyses of the GR interactome reveal a compositionally heterogeneous landscape that incorporates both coactivator (SWI/SNF, Mediator, EP300, NCOA2/3) and corepressor (SIN3A, NuRD, HDAC1/2) modules, with AR engaging an analogous but partly distinct repertoire (shared SWI/SNF and NuRD components plus an AR-selective NCoR/NuRD subnetwork). This suggests that the AR-GR complexes, and the condensates they nucleate, are compositionally flexible and can support both transcriptional activation and repression depending on chromatin context.

Mechanistically, AR and GR directly interact to form heterodimers on hormone response elements^24,25^. Although partial AR-GR genomic overlap has been observed in prostate cancer cell lines, direct biochemical evidence for heterodimerization was previously lacking. Co-IP of mouse prostate nuclear extracts, pull-down assays with recombinant proteins, and systematic deletion mapping collectively identify the GR LBD as the principal interface for AR binding, and structural modelling predicts an interface involving LBD helices H1 and H3, consistent with previous structural studies of the multidomain GR-DNA complex^50^. Direct heterodimer formation on DNA was confirmed by EMSA with extracts co-expressing AR and GR on probes derived from both the *Tmprss2* and *Ddit4* HREs, and by ChIP-reChIP at the *Ddit4* locus, demonstrating that both receptors co-occupy the same element *in vivo*. Functionally, this heterodimerization yields context-dependent outcomes: AR and GR may cooperate at *Tmprss2*-type (activating) HREs, but may also dampen each other at *Ddit4*-type (repressive) HREs, supporting a model in which HRE sequence composition and enhancer architecture help specify the functional output of AR-GR complexes.

Our genome-wide analyses further demonstrate extensive AR-GR co-occupancy of enhancer and super-enhancer regions, including at key lineage genes, in agreement with macromolecular condensate formation. This is consistent with tissue-culture studies in which AR and GR share 25-45 % of their binding sites depending on the cell type, with roughly 50 % of genes bound individually by either receptor regulated by both^1,23^. Although steroid receptors are known to converge on super-enhancers to stabilize lineage-specific programs^54,55^, AR-GR co-binding at these clusters had not been described in prostate under physiological conditions. Critically, genes associated with AR-GR co-bound enhancers can be either activated or repressed, indicating that GR does not merely repress AR-dependent transcription, as proposed^24^, but also actively promotes androgen-driven programs. Our RIME analyses provide a molecular basis for this duality: AR and GR converge on common activating machineries (SWI/SNF and Mediator), in line with independent RIME analyses in U2OS cells showing preferential GR recruitment of Mediator^56^, yet engage partially distinct corepressor modules, GR predominantly with the SIN3A/NuRD complex and AR predominantly with the NCoR/NuRD complex, both converging on shared HDAC1/2 activity. Consistent with this, more than 90% of AR- and GR-bound genes carry the H3K4me2 promoter/enhancer hallmark. Because H3K4me2-positive co-bound genes can be either activated or repressed, this mark is best interpreted as a readout of chromatin competence rather than of transcriptional direction, which is instead set by the cofactor complexes recruited to each locus. The identification of TP63, a master regulator of epithelial lineage programs^57,58^, among the most significantly enriched GR-associated transcription factors further positions AR-GR complexes within the hierarchy that governs epithelial identity.

A critical mechanistic finding is the identification of distinct DNA-sequence determinants that specify the output of AR-GR complexes. *De novo* motif discovery at AR-GR co-bound peaks associated with genes differentially expressed upon receptor silencing showed that activated targets (down-regulated upon silencing) are enriched for one HRE variant (5’-GGWACA*btb*TGTTCT-3’, matching the *Tmprss2* element), whereas AR-GR-repressed targets (up-regulated upon silencing) harbour a divergent variant (5’-RGAACA*dsh*TGTTCY-3’, matching the *Ddit4* element). This dichotomy offers a molecular basis for the context-dependent behaviour of AR-GR heterodimers. Subtle differences in half-site composition and spacer sequence within a shared HRE may bias whether coactivator- or corepressor-loaded complexes are recruited, reconciling the paradox that co-bound genes display uniformly permissive H3K4me2 yet opposing transcriptional outcomes. Together with the dual coactivator/corepressor architecture revealed by RIME and with the opposing behaviour of the *Tmprss2* (activating) and *Ddit4* (repressing) luciferase reporters, these findings define an “enhancer grammar” in which the HRE sequence contributes to selecting the mode of transcriptional regulation within a shared AR-GR framework.

The physiological relevance of these mechanisms is underscored by the prostate-specific GR knockout. GR loss in luminal cells disrupts AR nuclear condensation, producing a diffuse AR distribution, impairs expression of AR-dependent epithelial identity genes and de-represses nuclear-encoded mitochondrial and metabolic genes, phenocopying aspects of AR inhibition and indicating that GR is required for proper AR-driven differentiation *in vivo*. AR or GR silencing in organoids yields highly similar transcriptomes, confirming that the two receptors co-regulate a common gene network. Integrating the *in vivo* transcriptome with the GR cistrome shows that more than half of the genes dysregulated in GR^(i)pe-/-^ prostates are direct GR targets, with metabolic and translational programs (ribosome, oxidative phosphorylation, proteasome, mitophagy) de-repressed upon GR loss and epithelial signalling programs (adherens junction, focal adhesion, Wnt, MAPK, PI3K-Akt) down-regulated. A comparable fraction of these genes is also bound by AR, and these AR-bound gene recover the same metabolic and epithelial categories, indicating that the two receptors act on a largely shared set of genomic targets. This de-repression of a broad metabolic program is consistent with the NuRD, SIN3A, and NCoR/HDAC corepressor modules recovered in the AR-GR interactome, supporting a model in which GR normally restrains these genes through HDAC-containing complexes. At the tissue level, GR loss reduces prostate weight and glandular area, and elevates OXPHOS complex protein levels, whereas AR- or GR-silenced cells show increased mitochondrial respiration and glycolysis, indicating that AR-GR cooperation is required to maintain prostate epithelial homeostasis. Together, these findings support a model in which AR and GR act not as independent transcription factors but as interdependent partners within shared regulatory complexes. Ligand-activated GR promotes the assembly and dynamics of AR condensates. The two receptors heterodimerize through the GR LBD and co-occupy a common set of HREs at enhancer and super-enhancers. The local sequence of these elements, together with coactivator or corepressor complexes, determines whether AR-GR complexes activate or repress transcription. In this way, AR-GR cooperation ensures the precise execution of prostate epithelial transcriptional programs. More broadly, AR-GR cooperativity emerges as a fundamental property of normal prostate epithelial cell biology and a potentially actionable vulnerability in therapy-resistant prostate cancer.

## Materials and Methods

### Mouse Studies

#### Housing

All mice were maintained on a C57BL/6 background and housed in a temperature- and humidity-controlled facility under a 12-hour light/dark cycle, with *ad libitum* access to standard rodent chow (2800 kcal/kg, Usine d’Alimentation Rationelle, Villemoisson-sur-Orge, France) and water. Breeding and maintenance followed institutional guidelines. All experiments were performed in compliance with French and European Union regulations (Directive 2010/63/EU) and were approved by the local Ethical committee (Com’Eth, Strasbourg, France; APAFIS #37475).

Mice were euthanized by cervical dislocation, and prostates were weighed and collected for molecular and histological analysis.

#### Generation of GR^(i)pe-/-^ mutant mice

To selectively ablate GR in luminal prostate cells (GR^(i)pe-/-^), GR^L2/L2^ mice carrying LoxP sites flanking exon 3 encoding part of the GR DNA binding domain^48^ were intercrossed with PSA-CreER^T2^ mice that express the CreER^T2^ recombinase selectively in luminal prostate cells^51^. To induce recombination, eight-week-old GR^L2/L2^ control males and their PSA-CreER^T2^/GR^L2/L2^ pre-mutant littermates received intraperitoneal tamoxifen injections (1 mg/mouse/day) for five consecutive days, generating control and GR^(i)pe-/-^mutant mice. Primer sequences required for genotyping PCR are listed in Table S5-1.

#### Prostate histological analyses

Prostates were either snap-frozen in liquid nitrogen-cooled isopentane for cryosections, or fixed in 4% paraformaldehyde (PFA) overnight at 4 °C then washed in PBS and 70% ethanol and embedded in paraffin. 5 μm thick cryosections were prepared at -20 °C using a 2800 Frigocut cryostat (Leica, St Jouarre, France) and kept at -20 °C until further analysis. 5 μm thick paraffin sections were cut at room temperature using a microtome (Leica RM, 2145, 0737/09.1998) and stored at 4 °C until further analysis. Haematoxylin and eosin (H&E) staining was performed on paraffin sections as described^48^. Whole-prostate sections were imaged on a slide scanner (Digital Slide Scanner ZEISS Axioscan 7). For gland morphometry, H&E-stained sections of the dorsolateral prostate (DLP) were imaged and the mean glandular (luminal) area was quantified in QuPath.

For immunofluorescence, paraffin sections were dewaxed and subjected to antigen retrieval under controlled temperature and pressure conditions in 10 mM sodium citrate (pH 6.0, 20 min). Sections were permeabilized with 0.3% Triton X-100 in PBS, blocked with 5% BSA, and incubated overnight at 4 °C with primary antibodies directed against AR (Rabbit, Abcam, ab108341, 1:500), GR (Mouse, Santa Cruz Biotechnology, sc-393232, 1:200), CK18 (Mouse, Santa Cruz Biotechnology, sc-6259, 1:200). Mouse (Santa Cruz Biotechnology, E3117, 1:500), rat (Vector Laboratories, BA-4000, 1:500) or rabbit (PeproTech, 500-P000-500UG, 1:500) IgGs were used as controls. Alexa Fluor-conjugated secondaries (1:500, Invitrogen) were applied (1 hour, RT) and slides mounted in Fluoromount™ Aqueous Mounting medium with DAPI (Invitrogen, 00-4959-52).

For immunohistochemistry, mouse prostate cryosections were fixed in 4% PFA for 10 min. Endogenous peroxidase activity was quenched in 3% H_2_O_2_ in PBS for 10 min, and sections were blocked with 10% foetal bovine serum in PBS for 1 hour at room temperature. Primary antibodies directed against GR (Proteintech 24050, 1:200), AR (ab108341, 1:400) or anti-NKX3.1 (Proteintech 13069, 1:500), diluted in 0.5% BSA in PBS, were applied overnight at 4 °C. Slides were incubated with an HRP-conjugated anti-rabbit secondary reagent (SignalStain Boost IHC Detection Reagent, Cell Signaling Technology) for 1 hour at room temperature. Staining was developed with DAB substrate (SignalStain DAB Substrate, Cell Signaling Technology 8059) for 3 to 6 min and stopped with water. Sections were counterstained with Harris haematoxylin and mounted with Pertex.

### Prostate organoids

#### Prostate organoid isolation and culture

Mouse prostate epithelial organoids were generated from wild-type mice as previously described^59^. In brief, prostate was minced and digested with collagenase type II (5 mg/mL, Gibco) and dispase (1 mg/mL, Gibco) in Advanced DMEM/F12 (Invitrogen) supplemented with B27 (1x), 1.25 mM N-acetylcysteine, 50 ng/mL murine EGF, 100 ng/mL murine Noggin, 500 ng/mL murine R-spondin-1, 10 nM DHT, 10 μM Y-27632, and penicillin-streptomycin (100 U/mL) for 1 hour at 37°C with gentle agitation. Following digestion, tissue fragments were further dissociated with TrypLE (Gibco) for 10 min at 37 °C. Dissociated cells were filtered through a 70 μm cell strainer, centrifuged at 300 g for 5 min, and resuspended in ice-cold Matrigel (Corning, growth factor reduced). Cell-Matrigel suspension (40 μL droplets containing ∼5000 cells) was plated in pre-warmed 24-well plates and allowed to solidify for 20 min at 37°C. Medium was refreshed every 2-3 days. Organoids were passaged weekly at a 1:3-1:5 ratio using gentle mechanical dissociation after TrypLE treatment for 8 min.

#### Lentiviral production and organoid transduction

Lentiviral vectors encoding shRNAs targeting AR (shAR), GR (shGR), or a non-targeting control (shCtrl) in an EZ-Tet-pLKO-Puro vector backbone were produced in Lenti-X 293T cells (Takara Bio) using a third-generation packaging system^60,61^. Lenti-X 293T cells were co-transfected with psPAX2 packaging plasmid, pMD2.G envelope plasmid, and transfer vector using FuGENE HD (Promega) at a 3:1 ratio. Viral supernatants were collected 48-72 hours post-transfection, filtered through a 0.45 μm PVDF membrane, and concentrated using PEG-it (System Biosciences).

Organoids were dissociated to single cells using TrypLE supplemented with 10 μM Y-27632. Cells (1.5x10^4^ per condition) were incubated with concentrated lentivirus (MOI 5-10) in suspension for 30 min at room temperature in minimal volume (40 μL). Following incubation, cells were mixed with 120 μL cold Matrigel and plated as 40 μL droplets. Transduction efficiency was assessed by GFP expression at 72 hours post-transduction. Doxycycline (100 ng/mL) was added from day 4 post-transduction to induce shRNA expression.

### Cell culture

#### Cell lines and culture conditions

RWPE-1 human prostate epithelial cells (ATCC CRL-11609) were cultured in Keratinocyte-Serum-Free Medium (K-SFM, Gibco) supplemented with human EGF (5 ng/mL), bovine pituitary extract (0.05 mg/mL) and 10 nM dehydroepiandrosterone (DHEA). HEK293T (ATCC CRL-3216) and HeLa cells (ATCC CCL-2) were cultured in DMEM (Gibco) supplemented with 10% heat-inactivated foetal bovine serum, 2 mM L-glutamine, and penicillin/streptomycin (100 U/mL). Mammalian cells were kept at 37 °C, 5% CO₂. Sf21 cells (Thermo Fisher 11497013) were maintained in Grace’s Insect Medium supplemented with 10% FBS at 27 °C.

#### Cell treatments and transfections

For hormone treatments, RWPE-1 cells were cultured to 80% confluence, placed in hormone-deprived medium (K-SFM without supplements) for 24 hours, and treated with 10 nM DHT and/or 10 nM DEX (or 0.1% ethanol vehicle) for 24 hours.

For siRNA-mediated knockdown, RWPE-1 cells were transfected at 50-60% confluency using Lipofectamine RNAiMAX (Thermo Fisher, 13778075) according to the manufacturer’s reverse transfection protocol. ON-TARGETplus siRNA pools (Dharmacon L-003422-00-0005) targeting human *NR3C1* (GR) or non-targeting control siRNA were used at a final concentration of 20 nM.

For plasmid transfection, plasmids were introduced into HEK293T or HeLa cells with Lipofectamine 2000 (Thermo Fisher, 11668019) and cells were harvested 24 hours post-transfection.

#### Immunocytofluorescence (cells and organoids)

RWPE-1 cells grown on coverslips were fixed with 4% PFA in PBS for 15 min at room temperature, permeabilized with 0.5% Triton X-100 for 10 min, and blocked with 3% BSA in PBS for 30 min.

Organoids were fixed *in situ* with 4% paraformaldehyde in PHEM buffer (60 mM PIPES, 25 mM HEPES, 10 mM EGTA, 4 mM MgSO_4_, pH 6.9) for 30 min at RT. After quenching with 50 mM NH₄Cl, samples were permeabilized with 0.5% Triton X-100 in PBS for 30 min, and blocked with 5% BSA in PBS for 2 hours at 37 °C. Anti-AR (Rabbit, Abcam ab108341, 1:500) and anti-GR (Mouse, Santa Cruz, sc-393232, 1:200) were incubated overnight at 4 °C, followed by Alexa Fluor 488/594 secondaries (1:500, 2 hours, 37°C). Samples were mounted with DAPI and imaged on a Leica SP8 confocal (zoom 63x in oil). For condensate disruption experiments, cells were treated with 10% 1,6-HEX for 1 min (RWPE-1 cells) or 5 min (prostate organoids) at 37 °C immediately prior to fixation. For signal quantification, nuclear foci were analysed using ImageJ/FIJI software. Foci number and intensity were measured by applying a consistent threshold across all images within each experiment, and fluorescence intensity profiles were generated along linear regions of interest (ROIs) drawn across nuclei. Colocalization of AR and GR signals was assessed using the BIOP-JACoP plugin for ImageJ, and Manders’ colocalization coefficients (M1 and M2) were calculated from at least 50 cells per condition across three independent experiments^62^.

#### Single-molecule localization microscopy (SMLM)

For super-resolution imaging, RWPE-1 cells grown on 35 mm glass-bottom dishes fitted with a 14 mm #1.5 micro-well coverslip (Cellvis) were fixed in 4% PFA, immunostained with AR (Rabbit, Abcam ab108341, 1:200) and GR (Mouse, Santa Cruz sc-393232, 1:200) antibodies, as described above for immunofluorescence, and incubated with photoswitchable secondary antibodies CF680-conjugated anti-rabbit and AF647-conjugated anti-mouse, respectively. Two-colour acquisition used spectral-demixing single-molecule localization microscopy (splitSMLM)^63,64^. Imaging was performed in GLOX oxygen-scavenging buffer (200 U/mL glucose oxidase, 1000 U/mL catalase, 10% glucose, 200 mM Tris-HCl pH 8.0, 10 mM NaCl) supplemented with 50 mM MEA (cysteamine) and 2 mM COT (cyclooctatetraene) for two-colour direct stochastic optical reconstruction microscopy (dSTORM). Images were acquired with a modified Leica SR GSD system fitted with an HCX PL APO 160x/1.43 Oil CORR TIRF PIFOC objective. Fluorophores were first driven into the dark state with a 642 nm 500 mW fibre laser during an initial pumping phase in which emission was not recorded. Collection began once the density of emitting molecules was low enough to localize single fluorophores, and a 405 nm diode laser was applied as required to maintain an appropriate emitter density (back-pumping). Emission was relayed through an Optosplit II image splitter (Cairn Research) carrying a Chroma T690LPXXR dichroic mirror, generating a long- and a short-wavelength channel on an Andor iXon Ultra 897 EMCCD camera. The intensity ratio between the two channels was used to assign each localization to its fluorophore species (spectral demixing).

Single molecules were localized in both channels in Leica LAS X using the “direct fit” estimator. Localization tables were exported and processed with SharpViSu and the SplitViSu plug-in^63^ for spectral demixing, chromatic-aberration and drift correction, and image reconstruction. Super-resolution images were rendered as two-dimensional histograms of single-molecule coordinates at a 10 nm pixel size, and the magnified pore regions shown in the figures were upsampled by bilinear interpolation for display only. Colocalization of AR and GR on the reconstructed images was quantified with the BIOP-JACoP plugin^62^ (Manders’ M1/M2).

#### Subcellular fractionation

Cells were washed with cold PBS and resuspended in cytoplasmic extraction buffer (10 mM HEPES pH 7.6, 60 mM KCl, 1 mM EDTA, 0.075% NP-40, 1 mM DTT, 1 mM PMSF, and protease inhibitor cocktail). After 8 min on ice, samples were centrifuged at 400 g for 5 min and the supernatant was collected as the cytoplasmic fraction. Nuclear pellets were washed with PBS and resuspended in nuclear extraction buffer (20 mM Tris-HCl pH 8.0, 420 mM NaCl, 1.5 mM MgCl₂, 0.2 mM EDTA, 25% glycerol, 1 mM PMSF, and protease inhibitor cocktail). After 10 min on ice with occasional vortexing, samples were centrifuged at 15,000 x g for 10 min and the supernatant was collected as the nuclear fraction.

### Co-immunoprecipitation and western blotting

Mouse prostate tissue or cultured cells were lysed and nuclear extracts prepared by subcellular fractionation. Nuclear protein (500 μg) was incubated with 30 μL Protein G Dynabeads (Thermo Fisher Scientific, 10004D) and 5 μg of anti-AR (Abcam ab108341), anti-GR (IGBMC #3249), TRP63 (Santa Cruz sc8343) or rabbit IgG control antibody (Santa Cruz sc2357) in IP buffer (50 mM Tris-HCl pH 8.0, 170 mM NaCl, 0.1% NP-40, 20% glycerol, 50 mM NaF, 2 mM sodium orthovanadate, 0.2 mM DTT, PMSF, and protease inhibitor cocktail) for 2 hours at 4 °C with rotation. Beads were washed three times with cold IP buffer, and bound proteins were eluted by boiling in 2x Laemmli buffer for 5 min. The loading volumes on the gel were adapted to optimize interaction visualization. In brief, the 40 µL obtained after IP of Ab1 was loaded as 10 µL for western blot detection with Ab1 and 30 µL for western blot detection with Ab2. Proteins were resolved by SDS-PAGE, transferred onto PVDF membranes, and probed using antibodies against AR (Abcam ab108341, 1:1000), GR (IGBMC #3249, 1:1000), TRP63 (Santa Cruz sc8343) or α-Tubulin (1:5000), followed by HRP-conjugated secondary antibodies. Immunoreactive bands were detected using enhanced chemiluminescence substrate (Pierce) and visualized on an ImageQuant800 imaging system (Cytiva). Band intensities were quantified using Fiji/ImageJ.

For immunoprecipitation experiments performed in transfected HEK293T cells expressing Flag-tagged proteins, anti-Flag M2 affinity gel (Sigma F1804) was used for direct capture of tagged proteins.

### Rapid immunoprecipitation mass spectrometry (RIME)

RIME was performed on RWPE-1 cells as previously described^46^. Cells were treated with 100 nM DHT and 100 nM DEX for 2 hours, and then cross-linked by adding 1% formaldehyde (EM grade, methanol-free) directly to the culture medium for 10 minutes at room temperature, followed by quenching with glycine. Sonicated lysates were incubated overnight at 4 °C with 10 μg of anti-AR, anti-GR antibodies, or rabbit IgG control. Immune complexes were captured using Protein G Dynabeads. Beads were washed 10 times with cold RIPA buffer, followed by washes with cold 100 mM ammonium bicarbonate. Proteins were subjected to on-bead tryptic digestion overnight. Digested peptides were desalted and analysed by LC-MS/MS on an Orbitrap Fusion mass spectrometer (Thermo Fisher Scientific). Raw data were processed using MaxQuant software against the human UniProt database. Differential enrichment of each bait over the matched IgG control was assessed from three biological replicates using a two-sided Student’s t-test on log₂ LFQ intensities (fold-change and p/FDR cutoff). Enriched interactors were displayed as volcano plots. Protein-protein interaction networks among the significantly enriched proteins were constructed in STRING (v12.0)^65^ at a medium-confidence score.

### Chromatin Immunoprecipitation experiments

#### Chromatin Immunoprecipitation

ChIP was performed as previously described^48^ with minor modifications. Tissues were minced and cross-linked with 4% PFA in PBS for 10 min at room temperature with rotation, quenched with 125 mM glycine for 5 min, and washed with cold PBS. Cells were cross-linked directly in culture dishes. Cross-linked samples were lysed in ChIP lysis buffer with protease inhibitors and sonicated. Sonicated chromatin was diluted 10-fold in ChIP dilution buffer and pre-cleared with Protein G Dynabeads for 1 hour at 4 °C. For each ChIP, 250 μg of chromatin was incubated with 5 μg of antibody (anti-AR Abcam ab108341, anti-GR IGBMC #3249, or anti-H3K4me2 Active Motif #39141) overnight at 4°C with rotation. Immune complexes were captured with Protein G Dynabeads for 2 hours and washed sequentially with low-salt, high-salt, LiCl wash buffers, and TE buffer. Chromatin cross-links were reversed overnight at 65 °C with 200 mM NaCl and 20 μg RNase A. Samples were treated with proteinase K (50 μg) for 2 hours at 55°C, and DNA was purified using phenol-chloroform-isoamyl extraction followed by ethanol precipitation. For ChIP-qPCR, immunoprecipitated and input DNA were analysed by quantitative PCR using SYBR Green master mix on a LightCycler 480 II. Primers were designed to amplify 80-150 bp regions flanking predicted AR/GR binding sites (Table S5-2). Enrichment was calculated relative to input DNA using the percent input method.

#### Sequential ChIP (ChIP-reChIP)

For re-ChIP experiments to detect AR-GR co-occupancy, the first ChIP was performed as described above using anti-AR or anti-GR antibodies (anti-AR Abcam ab108341, anti-GR IGBMC #3249, or rabbit IgG control Santa Cruz sc2357). After washing, immune complexes were eluted with 10 mM DTT in TE buffer at 37 °C for 30 min instead of the standard elution buffer. The eluate was diluted 50-fold in ChIP dilution buffer and subjected to a second immunoprecipitation with anti-GR or anti-AR antibodies, respectively, or an IgG control overnight at 4 °C. The second ChIP was completed as described for standard ChIP, and purified DNA was analysed by qPCR. Primers are described in Table S5-2.

#### ChIP-sequencing

ChIP-seq libraries were prepared from 1-20 ng of purified DNA at the GenomEast platform using the Diagenode MicroPlex Library Preparation kit v3. Libraries were sequenced on an Illumina HiSeq 4000 sequencer as single-read 50 base reads. Image analysis and base calling were performed using RTA version 2.7.7 and bcl2fastq (version 2.20.0.422). Reads were mapped to the mm10 reference genome using Bowtie2 (version 2.4.4)^66^, and reads aligning to the ENCODE mm10 blacklist (V2) were filtered out. Peak calling was performed using MACS2 (version 2.2.7.1)^67^ with input DNA as control. All peaks with an FDR greater than 0.05 were excluded from further analysis. ChIP-seq was performed in two independent biological replicates per antibody. Peaks were annotated using HOMER software (v4.11.1) with mm10 Refseq annotation^68^. Genomic features (promoter/TSS, 5’ UTR, exon, intron, 3’ UTR, TTS and intergenic regions) were defined and calculated using Refseq and HOMER according to the distance to the nearest TSS. *De novo* motif discovery was performed using HOMER and the MEME Suite^69^, using 100 bp sequences extracted around peak summits. For bioinformatics and clustering analysis, Venn diagrams were generated using Venny (2.1.0)^70^. Clustering analyses were performed with the seqMINER software^71^, and deeptools^72^ (-computeMatrix -skip0, -plotHeatmap) with K-Mean linear normalization. Bigwig files were generated using bamCoverage (deeptools 3.3.0)^72^, with normalization by RPKM (--normalizeUsing RPKM --binSize 20). Genomic intersections and sequence extractions were carried out using bedtools: GetFastaBed for FASTA extraction, Intersect intervals, Multiple Intersect for overlapping locations, and WindowBed for gene-centric peak distribution. Super-enhancers were identified with the ROSE algorithm as described^41,42^. When not specified, all bioinformatics analyses were performed using default parameters.

### RNA extraction and analyses

Total RNA was isolated from tissues, cells, or organoids using TRIzol Reagent (Life Technologies Cat# 15596026) according to the manufacturer’s protocol. RNA concentration was measured using a NanoDrop spectrophotometer, and RNA integrity was assessed using an Agilent 2100 Bioanalyzer. Only samples with RNA Integrity Number (RIN) ≥ 8.0 were used for RNA-seq.

For RT-qPCR, 1 μg of total RNA was treated with DNase I to remove genomic DNA contamination and reverse transcribed using SuperScript IV Reverse Transcriptase (Life Technologies Cat# 18090010) with random hexamer primers. Quantitative PCR was performed using SYBR Green Master Mix on a LightCycler 480 II instrument (Roche). All samples were analysed in triplicate, and relative expression levels were calculated using the 2^-ΔΔ*Ct*^ method with normalization to housekeeping genes. Primer sequences are provided in Table S5-3.

For RNA-seq, libraries were prepared from 400 ng RNA and sequenced on an Illumina NextSeq 2000 (paired-end 2 x 50 bp), following the manufacturer’s instructions. Base calling used RTA 2.7.7 and bcl2fastq 2.17.1.14. Raw sequencing reads were preprocessed using cutadapt (version 4.2) to remove adapter, polyA, and low-quality sequences (Phred score<20). Reads shorter than 40 bases were discarded. Remaining reads were mapped to rRNA sequences using bowtie2 (version 2.3.5), and mapping reads were removed. Filtered reads were mapped onto the mm10 genome (GRCm38 assembly) from *Mus musculus* using STAR (version 2.7.10b)^73^. Gene expression quantification was performed from uniquely aligned reads using STAR version 2.7.10b^73^ with annotations from Ensembl version 102. Only non-ambiguously assigned reads were retained. Read counts were normalized across samples using the median-of-ratios method. For comparison among datasets, the transcripts with more than 50 raw reads were considered. Differential gene expression analysis was performed using the DESeq2 package (version 1.34.0)^74^ with a p < 0.05. Log2 Fold-Change shrinkage was performed with the apeglm method. Identified DEGs were subjected to functional analysis using STRING 12.0^65^ and to pathway/ontology enrichment using WebGestalt^75^ (Over-Representation Analysis against Gene Ontology Biological Process and KEGG pathways, FDR < 0.05). Gene set enrichment analysis (GSEA) was performed on the ranked gene lists using WebGestalt GSEA. Heatmaps were generated by centring and normalizing expression values using Cluster 3.0^76^, with subsequent visualization in Morpheus (https://software.broadinstitute.org/morpheus/). Genes were clustered using K-Mean with gene tree construction, Pearson correlation, and average linkage^77^.

### Recombinant protein expression and purification

Bacterial expression: His_6_-tagged human full-length AR, GR, and deletion mutants were cloned into pET28a expression vectors and transformed into *E. coli* BL21(DE3) competent cells. Bacterial cultures were grown in LB medium with kanamycin (50 μg/mL) at 37 °C to an OD_600_ of 0.6-0.8, induced with 0.5 mM IPTG, and incubated at 18 °C overnight with shaking. Cells were harvested by centrifugation, resuspended in lysis buffer (50 mM Tris-HCl pH 7.5, 200 mM NaCl, 4 mM DTT, lysozyme, 0.75 μg/mL DNase I and RNase A, 1x protease inhibitor cocktail), and lysed by sonication on ice. The lysate was clarified by centrifugation at maximum speed for 30 min at 4°C, and the supernatant was applied to Ni-NTA agarose resin (Qiagen) pre-equilibrated in lysis buffer. The column was washed with wash buffer (50 mM Tris-HCl pH 7.5, 200 mM NaCl, 4 mM DTT, 2 mM β-mercaptoethanol, 1x protease inhibitor cocktail), and proteins were eluted with elution buffer (50 mM Tris-HCl pH 7.5, 200 mM NaCl, 4 mM DTT, 2 mM β-mercaptoethanol, 250 mM imidazole, 1x protease inhibitor cocktail). Eluted proteins were concentrated using Amicon Ultra centrifugal filters (Millipore). Protein concentration was determined by Bradford assay, and purity was assessed by SDS-PAGE followed by Coomassie blue staining.

For insect cell expression, recombinant human AR and GR were expressed in Sf21 cells using the baculovirus expression system. Cells were infected at 2x10^6^ cells/mL with appropriate MOI and cultured for 48-72 hours at 27 °C. For AR expression, 10 nM DHT was added to the culture medium. For GR, 10 nM dexamethasone was used. Cells were lysed in lysis buffer (50 mM Tris-HCl pH 7.5, 500 mM NaCl, 5% glycerol, 0.1% NP-40, 1 mM TCEP, 1 mM β-mercaptoethanol, protease inhibitor cocktail, 10 μM ligand) using sonication (30% amplitude, 2x30 seconds, 0.5 sec on/off), clarified by centrifugation (15,000 x g, 30 min, 4 °C), and purified using appropriate affinity resins (Ni-NTA for His-tagged proteins, amylose resin for MBP fusions).

For fluorescently tagged proteins for LLPS assays, recombinant GFP-tagged full-length AR (GFP-AR^FL^) and mCherry-tagged GR (mCherry-GR^FL^) or GR^ΔLBD^ were expressed in *E. coli* BL21(DE3) cells as MBP fusion proteins. Expression was performed in LB medium supplemented with 0.1% glucose, kanamycin (50 μg/mL), 0.1 mM IPTG, and appropriate ligands (10 μM DHT for AR constructs, 10 μM DEX for GR^FL^) at 16 °C overnight. Cell pellets were lysed in LLPS lysis buffer (20 mM Tris-HCl pH 7.5, 400 mM NaCl, 10% glycerol, 3 mM TCEP, 0.1% NP-40, 0.5 mM CHAPS, 2x protease inhibitor cocktail, benzonase nuclease, 5 mM MgCl₂), supplemented with 5 μM triamcinolone for GR^FL^ constructs or 1 μM DHT for AR constructs. Proteins were purified using amylose resin, eluted with 10 mM maltose, and dialysed into low-salt LLPS buffer (20 mM Tris-HCl pH 7.5, 150 mM NaCl, 1 mM DTT, 5% glycerol).

### Bacterial expression for GST pull-downs

For GST pull-down experiments, GST-tagged full-length human GR (GST-hGR), GST-tagged human GR ligand-binding domain (GST-hGR-LBD), and GST alone were expressed in *E. coli* BL21(DE3) cells. Cultures were induced with 1 mM IPTG at OD600 0.4-0.6 and incubated overnight at 18 °C in the presence of 10 μM dexamethasone. Cells were lysed by sonication in 25 mM Tris-HCl pH 7.5, 100 mM NaCl and 2 mM EDTA. Lysates were clarified by centrifugation (10,000 x g, 20 min, 4 °C) and GST fusion proteins purified using Glutathione Sepharose 4B resin (Sigma, 17-0756-01).

### In vitro MBP pull-down assay

Sf21 cell pellets co-expressing Avi-MBP-GR and Avi-AR were lysed in 50 mM Tris-HCl pH 7.5, 250 mM NaCl, 5% glycerol, 0.01% NP-40 (Igepal), 1 mM CHAPS, 1 mM TCEP, 1 mM β-mercaptoethanol supplemented with benzonase nuclease, 10 mM MgCl₂, protease inhibitor cocktail, 0.1 mM PMSF, and the corresponding ligands. Lysates were sonicated, clarified by centrifugation, and incubated with amylose resin for 1.5-2 hours at 4 °C. Beads were washed extensively and bound proteins were eluted in SDS sample buffer. Input, flow-through, and eluate fractions were analysed by SDS-PAGE followed by Coomassie staining or immunoblotting using anti-AR and anti-GR antibodies. AR expressed alone served as a control for nonspecific binding to the amylose resin.

### GST pull-down assay

GST-tagged full-length GR (GST-GR), GST-tagged GR ligand-binding domain (GST-GR-LBD), or GST alone immobilized on Glutathione Sepharose 4B resin were incubated with purified His-tagged AR for 3 hours at 4 °C. Beads were washed three times in PBS containing 1% Triton X-100 and bound proteins were analysed by immunoblotting using antibodies against AR, GR, or GST (Merck/Millipore 05-782).

### Electrophoretic mobility shift assay (EMSA)

HEK293T cells were transfected with 8 μg of AR or GR pSG5 expression vectors. Forty-eight hours after transfection, cells were treated with 100 nM dexamethasone and/or 100 nM DHT for 1 hour, and whole-cell extracts were prepared in lysis buffer containing 20 mM Tris-HCl pH 7.5, 1 mM EDTA, 2 mM DTT, 20% glycerol, and 400 mM KCl.

Cy5-labelled double-stranded oligonucleotide probes (Eurogentec) encompassing the *Ddit4* BSe2 HRE (core 5’-AGAACAttgTGTTCT-3’) or the *Tmprss2* HRE (core 5’-GGAACTcttTGTTCA-3’) were generated by annealing complementary strands. Binding reactions (15 μL) contained 25 μg whole-cell protein extract, 133 μg/mL poly(dI-dC), 100 nM ligand (DHT and/or DEX), and binding buffer (10 mM Tris-HCl pH 7.5, 0.5 mM EDTA, 0.5 mM DTT, 5% glycerol, protease inhibitors, and 1 mM β-mercaptoethanol). For reactions with the *Tmprss2* probe, 1 mM MgCl₂ was added to reduce non-specific binding. After incubation for 30 min at 4 °C, 200 fmol Cy5-labelled probe was added and reactions were incubated for a further 20 min at 37 °C. Protein-DNA complexes were resolved on a non-denaturing 2-8% gradient polyacrylamide gel cast in 0.5x TBE containing 2.5% glycerol. Cy5 fluorescence was detected using an ImageQuant imaging system (Cytiva).

### Fluorescence recovery after photobleaching

HeLa cells seeded on glass-bottom dishes (MatTek) were transfected with eGFP-AR and/or mCherry-GR expression vectors. Twenty-four hours post-transfection, cells were treated with 10 nM DHT and 10 nM DEX for 24 hours to induce nuclear condensate formation Live-cell imaging was performed on a Zeiss LSM780 confocal microscope equipped with a temperature-controlled chamber (37 °C, 5% CO2) and a 63x oil-immersion objective. Individual nuclear condensates were selected as regions of interest (ROI, typically 2-3 μm in diameter), and baseline fluorescence intensity was recorded for 10 frames (5 seconds). The ROI was photobleached using 100% laser power (488 nm for GFP, 561 nm for mCherry) for 10 iterations until fluorescence intensity dropped to 20-30% of the pre-bleach level. Fluorescence recovery was then monitored by acquiring images every 0.5 seconds for 60 seconds post-bleaching at low laser power (2-5%). Fluorescence intensities were measured using ImageJ, background-subtracted, and normalized to the pre-bleach intensity. Recovery curves were fitted to a single exponential equation: F(t) = F_∞_ (1 - e^(-t/τ)^), where F_∞_ is the mobile fraction and τ is the recovery half-time. At least 10-15 condensates from different cells were analysed for each condition.

### Semi-denaturing detergent agarose gel electrophoresis assay

The formation of polymer-like assemblies and higher-order oligomers of recombinant AR and GR was analysed by SDD-AGE as described^78^. Recombinant proteins (1-5 μM) were incubated under phase separation conditions (50 mM Tris-HCl pH 7.5, 150 mM NaCl, 10% PEG8000) for 30 min at room temperature. Samples were mixed with 4x sample buffer (2% SDS, 10% glycerol, 0.01% bromophenol blue in TAE) without boiling or reducing agents to preserve native oligomeric states. Samples were loaded onto vertical 1.5% agarose gels containing 0.1% SDS and electrophoresed at 100 V for 1.5-2 hours at 4 °C in running buffer (TAE with 0.1% SDS). Proteins were transferred to PVDF membranes (Millipore). Membranes were blocked and immunoblotted with primary antibodies against AR (Abcam ab108341, 1:1000) or GR (IGBMC #3249, 1:1000) overnight at 4 °C. After washing with TBS-T, membranes were incubated with HRP-conjugated secondary antibodies (1:5000) for 1 hour at room temperature. Immunoreactive bands were detected using enhanced chemiluminescence substrate (Pierce) and visualized on an ImageQuant800 imaging system (Cytiva). Monomeric, oligomeric, and high-molecular-weight assemblies were identified based on their migration patterns relative to protein molecular weight markers. For quantification, band intensities were measured using ImageJ/FIJI software, and the ratio of oligomeric to monomeric species was calculated for each condition.

### Liquid-liquid phase separation (LLPS) assay

For phase separation assays, purified fluorescently tagged proteins were mixed at indicated concentrations (typically 5 μM each) in LLPS assay buffer (50 mM Tris-HCl pH 7.5, 150 mM NaCl) supplemented with 10% polyethylene glycol 8000 (PEG8000) as a molecular crowding agent. Samples (5-10 μL) were immediately loaded onto glass-bottom dishes (Cellvis) and observed by confocal microscopy (Leica TCS SP8) using a 40x oil-immersion objective. Droplet formation was monitored in real-time, and images were captured at 1 min intervals for 30 min. For quantitative analysis, droplet size distribution and number were measured using ImageJ/FIJI software. To assess the liquid-like properties of condensates, fusion events were recorded by time-lapse imaging^79^.

### Luciferase bulk reporter assay

HEK293T cells were seeded in poly-D-Lysine coated black chimney-style 96-well plates at a density of 3 x 10⁵ cells/mL and co-transfected the same day in biological duplicate. Cells were transfected with eGFP as internal control (10 ng), *Tmprss2*- or *Ddit4*-responsive firefly luciferase pGL4 reporter plasmids (150 ng) and either GR alone (37.5 ng), AR alone (37.5 ng), or both GR and AR pSG5 expression plasmids at different GR: AR ratios: 1:1, 1:2, or 1:3 (ratio 1 corresponding to 37.5 ng), as indicated. Cells were treated with DMSO (vehicle), DEX (20 nM) and/or DHT (20 nM), as indicated for GR-, AR- or combined conditions. Cells were washed once with PBS. Fluorescence signal was measured in duplicate using a GloMax® 96-well plate reader (Promega). Subsequently, 50 μL per well of ONE-Glo™ EX reagent (Promega, E8110) was added and incubated for 10 min at room temperature according to the manufacturer’s instructions. Luminescence was then measured in duplicate using the same instrument, and relative luciferase units (RLU) were calculated as the ratio of luminescence to eGFP fluorescence.

### Seahorse metabolic flux analysis

Mitochondrial respiration and glycolytic activity were assessed using the Seahorse XF Cell Mito Stress Test Kit (Agilent Technologies, 103015-100) and the Seahorse XF Glycolysis Stress Test Kit (Agilent Technologies, 103020-100), according to the manufacturer’s protocols. Briefly, RWPE-1 cells transfected with siCtrl, siAR or siGR were seeded at 4 x 10⁴ in Seahorse XF cell culture microplates (Agilent). Extracellular acidification rate (ECAR) and oxygen consumption rate (OCR) were measured on a Seahorse XFe96 analyzer (Agilent). For the glycolysis stress test, glucose (10 mM), oligomycin (1 µM) and 2-deoxyglucose (50 mM) were injected sequentially, and glycolysis, glycolytic capacity and glycolytic reserve were calculated. For the mitochondrial stress test, oligomycin (1 µM), FCCP (2.5 µM) and rotenone/antimycin A (1 µM) were injected sequentially, and basal respiration, ATP-linked respiration, maximal respiration and spare respiratory capacity were calculated. Parameters were calculated with Wave software (Agilent) and normalized to nucleus number.

### Structure Modelling

The modelling was performed online using AlphaFold 3 (https://alphafoldserver.com/)^80^. The sequences of the various proteins were informed in the server: one copy of hGR LBD (aa 522-777), one copy of hAR LBD (aa 660-920) and one copy of an LxxLL-type coregulator box. Two different 50 amino acid long peptides were used, either the FxxLF-containing peptide of AR (1-50), or the LVQLL-containing peptide of NCOA2 (TIF2). AlphaFold Server then allowed modelling of the complex consisting of AR LBD, GR LBD and the peptide. The confidence score and the various models were then examined using PyMOL Molecular Graphics System, Version 3.1.6.1 Schrödinger, LLC.

### Statistical analysis

All data are presented as mean ± standard error of the mean (SEM) unless otherwise indicated. Biological replicates represent independent experiments performed on different days with independently prepared samples (different animals, different cell passages, or different organoid preparations). Statistical significance was determined using GraphPad Prism software (version 10). The specific statistical tests used (e.g., two-tailed Mann-Whitney test, two-way ANOVA with Tukey’s post hoc test) are indicated in the figure legends for each experiment. The number of biological replicates is indicated in the figure legends.

## Resource Availability

The raw and processed high-throughput sequencing datasets for RNA-seq and ChIP-seq have been deposited to the Gene Expression Omnibus (GEO) database under accession number GSE312593 (RNA-seq in mouse organoids), GSE330088 (RNA-seq in mouse prostate) and GSE312487 (ChIP-seq in mouse prostate), respectively. H3K4me1, H3K4me3 and H3K27ac datasets were taken from GSM1145319, GSM1145320 and GSM1259357, respectively. All remaining data are available in the Article, Supplementary, and Source Data files.

## Acknowledgments

We are grateful for the valuable technical support of Régis Lutzing (tamoxifen treatment, reagent ordering and laboratory management assistant), Anastasia Bannwarth, Nikola Djordjevic and Brayann Weis (genotyping and plasmid preparation), Paola Rossolillo (lentivirus production) and Mustapha Oulad-Abdelghani (antibody generation and purification). We are grateful to George Couch and Soleilmane Omarjee for critical reading. We thank Darya Yanushko and Noé Pierrat for training in prostate organoid isolation, culture and immunofluorescence, Kateryna Len for training in prostate histology, Valentine Gilbart and Céline Keime for bioinformatics assistance, Marc Ruff for advice on the *in vitro* purification conditions, Shanming Ji for advice on the LLPS and SDD-AGE assays and Leonid Andronov for the splitSMLM microscopy. We also thank Prof. Shigeaki Kato for providing the AR-floxed mice. We acknowledge the support of the IGBMC core facilities, including the Integrated Structural Biology and Photonic Microscopy platforms, Molecular biology and Baculovirus services, and PluriCell East cell culture and Histopathology services, as well as the IGBMC animal facility, Mouse Clinical Institute (ICS, Illkirch-Graffenstaden, France) and GenomEast facility, a member of the ‘France Génomique’ consortium (ANR-10-INBS-0009).

This work was supported by the Centre National de la Recherche Scientifique (CNRS) and the Institut National de la Santé et de la Recherche Médicale (Inserm), and by IdEx Unistra (ANR-10-IDEX-0002), SFRI-STRAT’US (ANR-20-SFRI-0012) and EUR IMCBio (ANR-17-EURE-0023), within the framework of the French Investments for the Future Programme. Additional support was provided by Inserm, CNRS, the University of Strasbourg, IGBMC, the Ligue Contre le Cancer (CCIR Est) and the Agence Nationale de la Recherche (ANR-10-BLAN-1108 and AR2GR ANR-16-CE11-0009-01). Q.C. was funded by the ANR (ANDROMETAMUS, ANR-24-CE14-0247-01) and T.S. by the Ministère de l’Enseignement Supérieur, de la Recherche et de l’Innovation.

## Author contributions

The study was conceived by D.D., D.M, and I.B. The research plan and the conceptualization of the results were based on discussions between Q.C., I.B., D.D. and D.M. Q.C. performed most of the experiments and analysed the data. T.S., F.C. and J.W.L. performed the EMSA and luciferase reporter assays that characterized the AR-GR heterodimer. D.R. and A-I.R. performed the ChIP-seq experiments. F.C. and S.S.C. generated the GR mouse line. K.E. carried out plasmid construction. E.E. and S.P. performed the RIME experiments. L.B., G.R. and A.B. performed the histological analyses of mouse tissues. T.Y. performed the bioinformatic analyses. J.R. performed the Seahorse metabolic flux assays. R.S. and E.C. performed cell culture experiments. N.B. and H.R.V. purified the recombinant proteins, with input from G.T., M.D. and E.M. D.D., I.B. and D.M. supervised the project, and Q.C., D.D., I.B. and D.M. took primary responsibility for writing the manuscript with input from all authors.

## Declaration of generative AI and AI-assisted technologies

None declared.

