## Supplementary Figures 1 to 6 for "Interdependent androgen and glucocorticoid receptor signalling shapes prostate epithelial homeostasis"

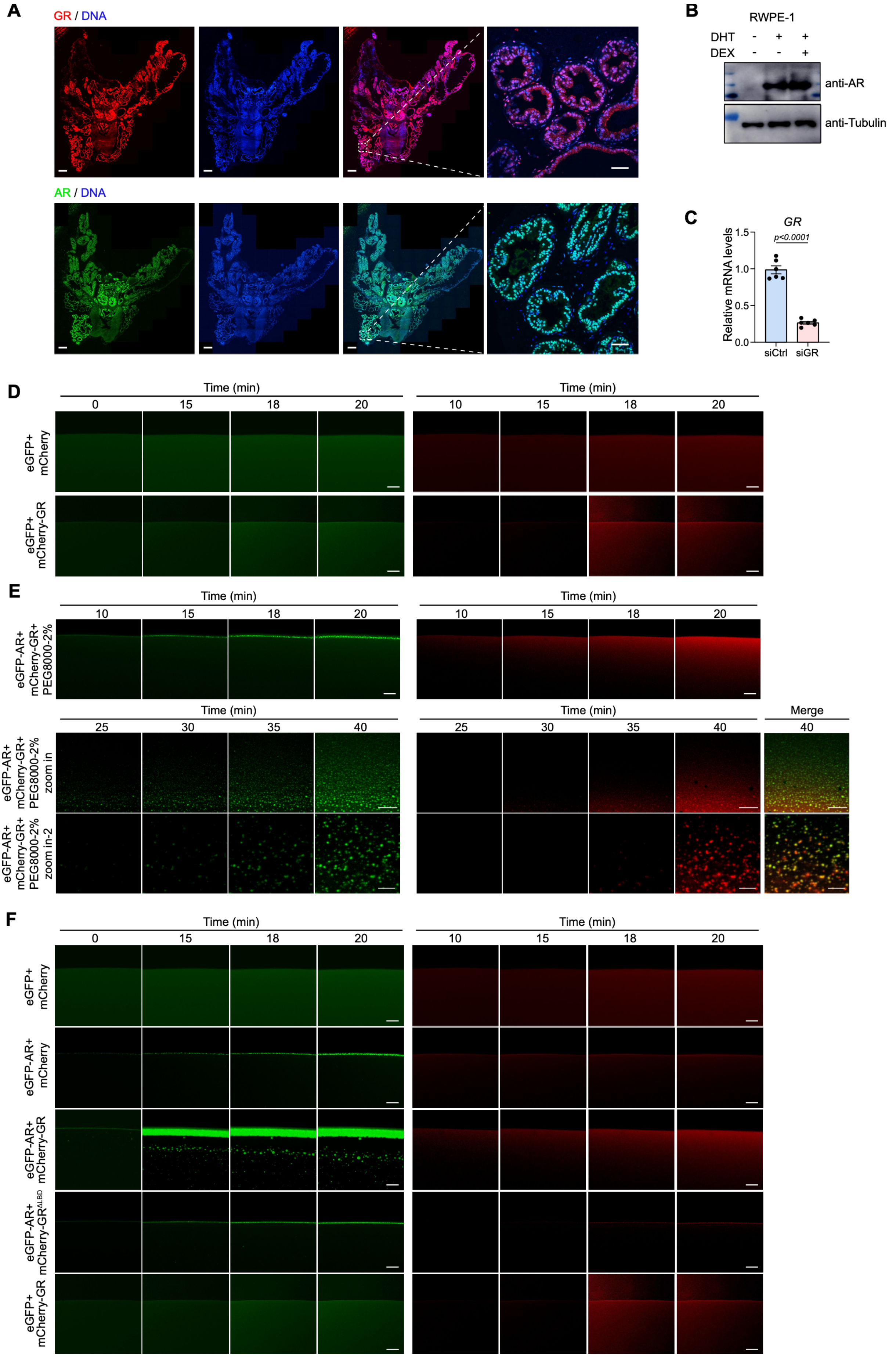
A
GR / DNA
AR / DNA
B
RWPE-1
DHT - + +
DEX - - +
anti-AR
anti-Tubulin
C
GR
p<0.0001
Relative mRNA levels
1.5
1.0
0.5
0.0
siCtrl
siGR
D
Time (min)
0
15
18
20
10
eGFP+ mCherry
eGFP+ mCherry-GR
E
eGFP-AR+ mCherry-GR+ PEG8000-2%
25
30
35
40
Merge
eGFP-AR+ mCherry-GR+ PEG8000-2% zoom in
eGFP-AR+ mCherry-GR+ PEG8000-2% zoom in-2
F
eGFP-AR+ mCherry
eGFP-AR+ mCherry-GR
eGFP-AR+ mCherry-GR$^{\Delta LBD}$

12

**Figure S1. AR and GR co-expression in prostate epithelium and characterization of LLPS assay components**

A. Immunofluorescence of GR (red, top) and AR (red, bottom) with DAPI (cyan) in adult mouse prostate sections. Left to right: single channel, DNA, merge, and a magnified view of the boxed region. Scale bar, 50 µm.

B. Immunoblot of AR in RWPE-1 cells treated with vehicle (-), DHT alone (+/-), or DHT + DEX (+/+). α-Tubulin serves as a loading control.

C. RT-qPCR of GR mRNA in RWPE-1 cells transfected with siGR or siCtrl. Mean ± SEM.

D. Time-lapse images of *in vitro* LLPS assays, eGFP + mCherry (top) and eGFP + mCherry-GR (bottom). Images at 0, 15, 18, and 20 min (left) and 10, 15, 18, and 20 min (right). Scale bar, 10 µm.

E. Time-lapse imaging of *in vitro* LLPS assays with eGFP-AR and mCherry-GR in 10% PEG8000. Top, overview (10-20 min). Bottom, two magnified regions showing droplet fusion at 25-40 min. Scale bar, 10 µm (top), 5 µm (bottom).

F. Time-lapse imaging of *in vitro* LLPS assays. Top to bottom: eGFP + mCherry, eGFP-AR + mCherry, eGFP-AR + mCherry-GR, eGFP + mCherry-GR, and eGFP-AR + mCherry-GR$^{\Delta LBD}$, at the indicated times (0-20 min). Scale bar, 10 µm.

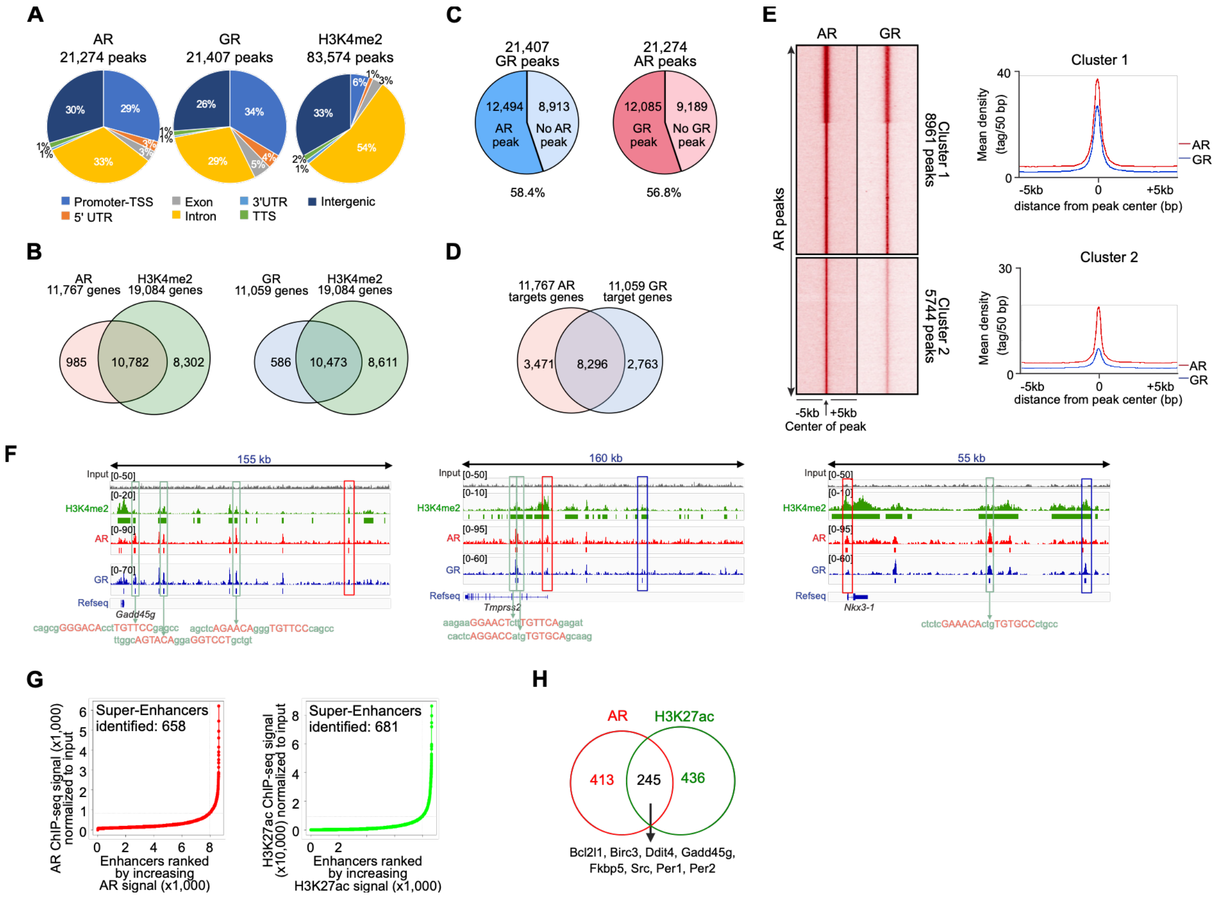


**Figure S2. Genome-wide characterization of AR and GR chromatin binding and super-enhancer identification**

A. Genomic distribution (pie charts) of AR, GR and H3K4me2 ChIP-seq peaks in mouse prostate.

B. Overlap (Venn) of genes bound by AR or GR and by H3K4me2.

C. Genomic distribution (pie charts) of AR- and GR-bound regions.

D. Overlap (Venn) of genes bound by AR and GR.

E. Tag-density heatmaps of AR and GR within ±5 kb of the AR peak centre, with average tag-density profiles and *de novo* motif analysis. The two first clusters were considered as Cluster 1.

F. Representative IGV tracks of AR, GR, H3K4me2, and Input ChIP-seq signals at *Gadd45g*, *Tmprss2*, and *Nkx3-1* loci. HRE sequences are indicated below each track.

G. Ranked enhancer-signal (hockey-stick) plots. 658 super-enhancers by AR ChIP-seq (left) and 681 by H3K27ac (right), ranked by increasing signal.

H. Overlap (Venn) of AR- and H3K27ac-defined super-enhancers. 245 shared, with representative genes *Bcl2l1*, *Birc3*, *Ddit4*, *Gadd45g*, *Fkbp5*, *Src*, *Per1*, and *Per2*.

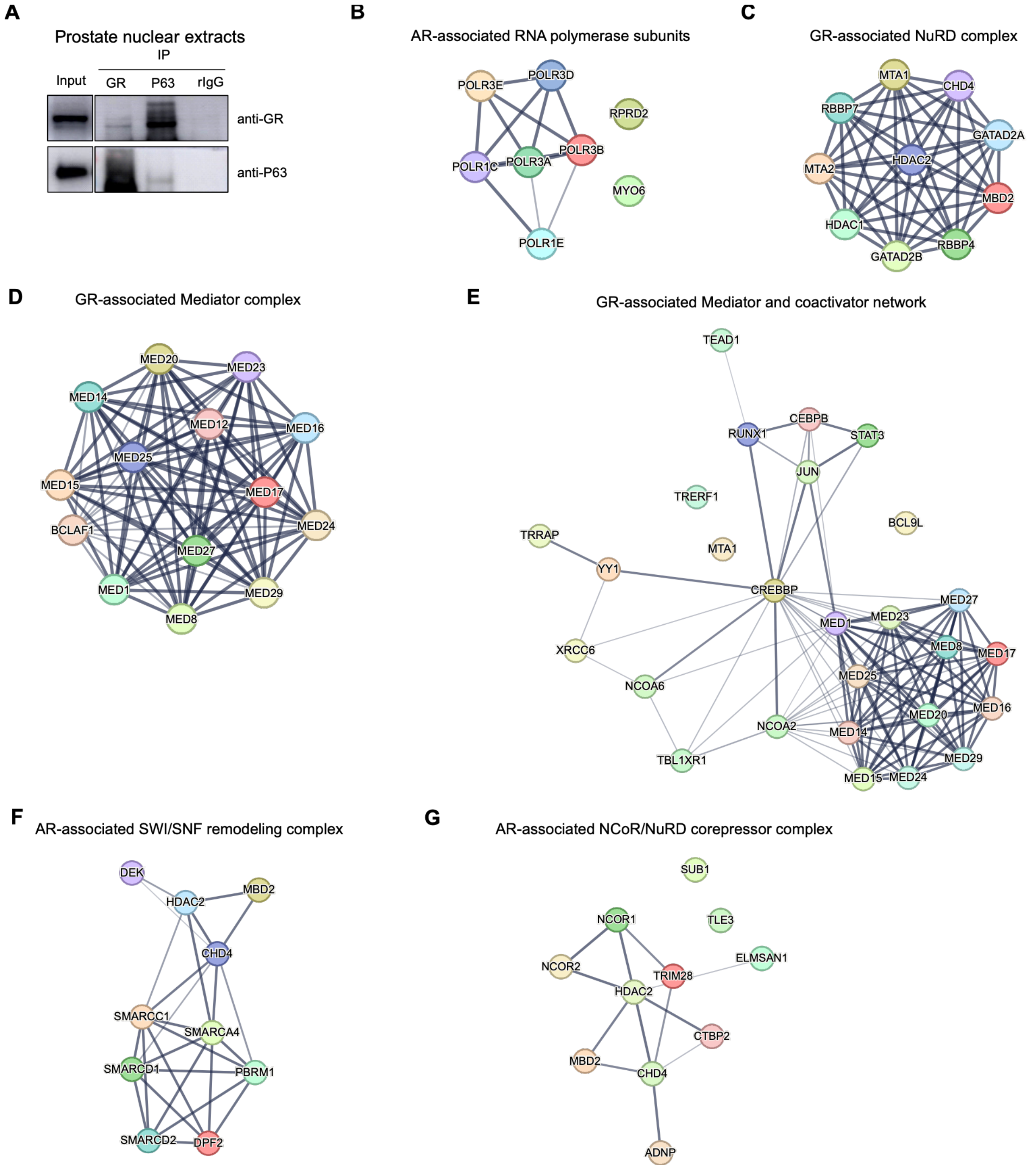


**Figure S3. AR- and GR-RIME interactomes and complex composition**

A. Co-immunoprecipitation of GR and TP63 from mouse prostate nuclear extracts (anti-GR, anti-TP63 or rabbit IgG. Immunoblotted for GR and TP63).

B. STRING network of AR-associated RNA polymerase module connected to the AR-RIME interactome.

C. STRING network of the GR-associated NuRD complex identified by RIME.

D-E. STRING network of GR-associated Mediator and coactivator components connected to the GR-RIME interactome: Mediator subunits (MED1, MED8, MED12, MED14-17, MED20, MED22-25, MED27, MED29) (D), coactivators (NCOA2, NCOA6, CREBBP, TBL1XR1, TRRAP) and associated factors (BCLAF1, JUN, RUNX1, STAT3, CEBPB, TEAD1, TRERF1, BCL9L, YY1, MTA1, XRCC6) (E).

F-G. STRING networks of the AR-associated SWI/SNF complex (F) and NCoR/NuRD corepressor complex (G).

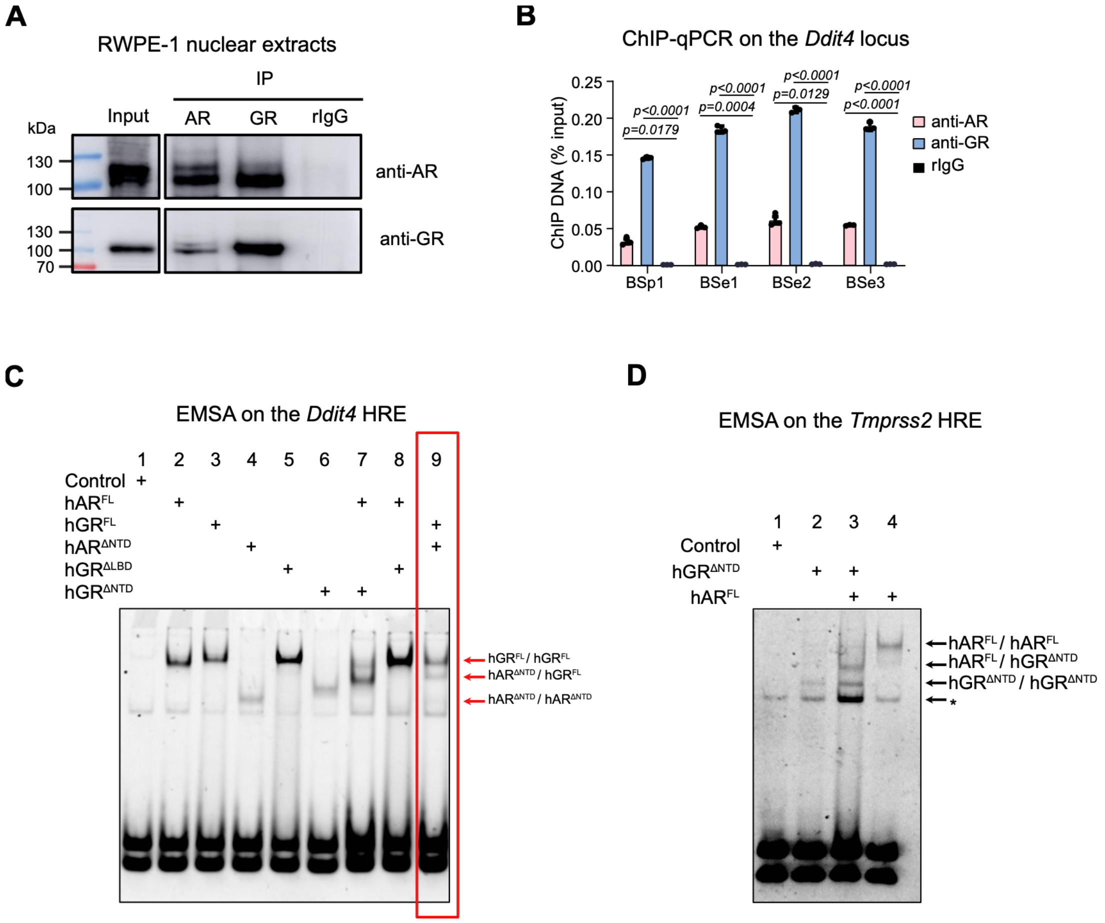


**Figure S4. Endogenous AR-GR interaction and response-element binding by AR and GR deletion mutants**

A. Co-immunoprecipitation of AR and GR from RWPE-1 nuclear extracts (anti-AR, anti-GR or rabbit IgG; immunoblotted for AR and GR).

B. ChIP-qPCR at the AR-GR binding sites (BS) of the *Ddit4* locus with the indicated antibodies. Mean ± SEM, one-way ANOVA with Tukey's correction.

C. EMSA on the BSe2 *Ddit4* response element with whole-cell extracts from HEK293T cells expressing the indicated AR or GR constructs. Lane 1, negative control. Lane 2, hAR$^{FL}$. Lane 3, hGR$^{FL}$. Lane 4, hAR$^{\Delta NTD}$. Lane 5, hGR$^{\Delta LBD}$. Lane 6, hGR$^{\Delta NTD}$. Lane 7, hAR$^{FL}$ + hGR$^{\Delta NTD}$. Lane 8, hAR$^{FL}$ + hGR$^{\Delta LBD}$. Lane 9, hGR$^{FL}$ + hAR$^{\Delta NTD}$.

D. EMSA on the *Tmprss2* HRE (5’-GGAACTcttTGTTCA-3’) with whole-cell extracts from HEK293T cells expressing Flag-hGR$^{\Delta NTD}$ and/or hAR$^{FL}$ in the presence of ligands. Lane 1, mock-transfected negative control. Lane 2, hGR$^{\Delta NTD}$. Lane 3, hGR$^{\Delta NTD}$ + hAR$^{FL}$. Lane 4, hAR$^{FL}$. Homodimeric (hGR$^{\Delta NTD}$/hGR$^{\Delta NTD}$, hAR$^{FL}$/hAR$^{FL}$) and heterodimeric (hAR$^{FL}$/hGR$^{\Delta NTD}$) complexes are indicated at right. Asterisks denote non-specific complexes present in all lanes, including the receptor-free control.

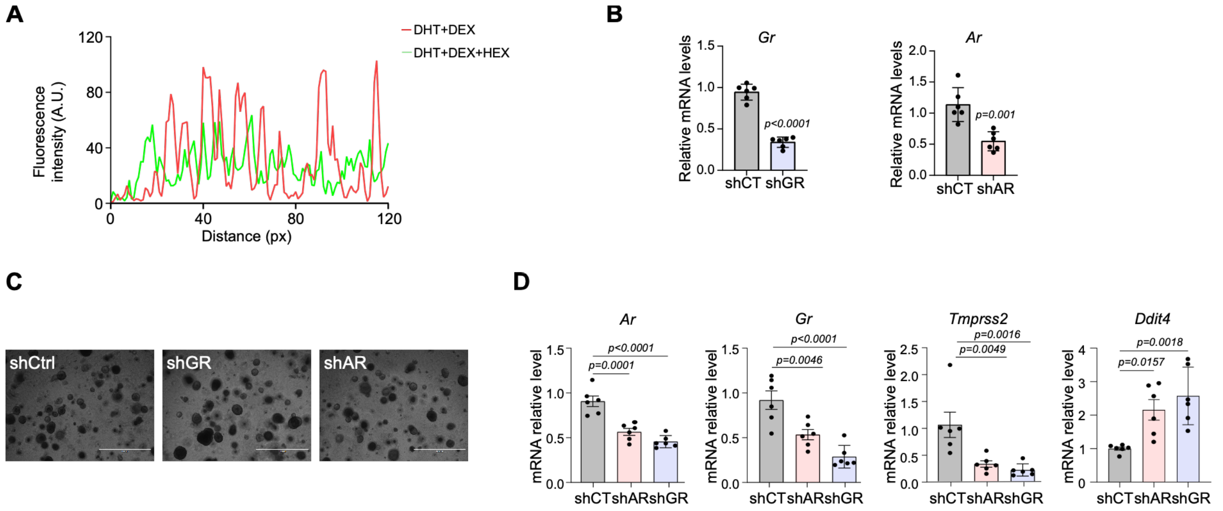


**Figure S5. AR nuclear condensation and validation of shGR and shAR prostate organoid models.**

A. Quantification of AR fluorescence intensity along line scans in prostate organoids treated with DEX (red) or with DEX and 10% 1,6-HEX (green).

B. RT-qPCR of GR and AR mRNA in prostate organoids transduced with shGR, shAR or control shRNA (shCtrl). Mean ± SEM.

C. Representative brightfield images of organoids transduced with shCtrl, shGR or shAR. Scale bar, 1,000 µm.

D. Relative transcript levels of the indicated genes in organoids transduced with shCtrl, shAR or shGR. Mean ± SEM. one-way ANOVA with Tukey's correction.

**Figure S6. Generation of prostate-specific GR knockout mice and metabolic phenotyping**

A. Strategy to generate tamoxifen-inducible, prostate-epithelium-specific GR knockout mice (GR$^{(i)pe-/-}$). PSA-CreER$^{T2}$ mice were crossed with *GR*$^{L2/L2}$ (floxed) mice. Tamoxifen-induced Cre excises exon 3 of the *Nr3c1* gene, generating a null allele.

B. PCR genotyping of control and GR$^{(i)pe-/-}$ mice. Cre transgene (primers p1-p2, 300 bp, top) and the recombined GR$^{L-}$ allele (primers p4-p5, 400 bp, bottom).

C. Immunofluorescence of GR (red) and DNA (DAPI, cyan) in prostate sections from control and GR$^{(i)pe-/-}$ mice. Scale bar, 1 mm.

D. Quantification of AR fluorescence intensity in prostate epithelial cells of control (red) and GR$^{(i)pe-/-}$ (green) mice. Mean ± SEM.

E. KEGG enrichment of genes up-regulated (left, pink) and down-regulated (right, blue) in GR$^{(i)pe-/-}$ prostate versus control prostate.

F. Comparison of the GR$^{(i)pe-/-}$ prostate and GR-silenced organoid transcriptomes. Overlap (Venn) of down-regulated DEGs (top; GR$^{(i)pe-/-}$ prostate, 3,604 and shGR organoids, 1,026; 391 shared) and of up-regulated DEGs (bottom; GR$^{(i)pe-/-}$ prostate, 2,284 and shGR organoids, 1,006; 425 shared), each shown with GO enrichment of the shared genes (right).

G. Integration of the AR cistrome and the GR$^{(i)pe-/-}$ transcriptome. Overlap (Venn) of up-regulated (2,284) and down-regulated (3,604) DEGs in GR$^{(i)pe-/-}$ prostate with AR ChIP-seq peaks (AR ChIP, 11,767), shown with KEGG enrichment of the AR-bound down-regulated (blue) and up-regulated (pink) genes.

H. Overlap of the AR and GR cistromes at genes dysregulated in GR$^{(i)pe-/-}$ prostate. Three-way overlap (Venn) of AR ChIP (11,767) and GR ChIP (11,059) peaks with down-regulated DEGs (left, 3,604) and with up-regulated DEGs (right, 2,284), each shown with KEGG enrichment of the genes co-bound by AR and GR.

I. Representative H&E-stained whole-prostate sections of control and GR$^{(i)pe-/-}$ mice. Scale bar, 1mm.

J. Seahorse assays in RWPE-1cells transfected with siCtrl, siAR or siGR. Top, extracellular acidification rate (ECAR, mpH/min) with sequential glucose, oligomycin and 2-DG injections (glycolysis, glycolytic capacity, glycolytic reserve and non-glycolytic acidification). Bottom, oxygen consumption rate (OCR, pmol $O_2$/min) with sequential oligomycin, FCCP and rotenone plus antimycin A injections (basal respiration, ATP-linked respiration, maximal respiration, spare respiratory capacity and non-mitochondrial respiration). Mean ± SEM.
